# Lysosomal Glycosaminoglycan Catabolism Regulates Microglial Cell State in the Developing CNS

**DOI:** 10.64898/2026.10.11.758365

**Authors:** Keenan Hope, Yiwen Zhang, Amrita Basu, Bingmin Luo, Kaleb Chassie, Ryan J. Weiss, Dhananjay Yellajoshyula

## Abstract

Microglia acquire their adult homeostatic identity during postnatal CNS development, a period of extensive extracellular matrix (ECM) remodeling, through mechanisms not fully understood. Here we show that lysosomal catabolism of glycosaminoglycans (GAGs), major ECM components, is a microglial function required to establish homeostatic identity. Using mouse models of Mucopolysaccharidosis VII, we examined the consequences of deleting β-glucuronidase (GUSB), a lysosomal enzyme required for GAG degradation. Loss of GUSB resulted in the accumulation of undegraded chondroitin sulfate within microglial lysosomes, confirmed by LC/MS analyses of cortical microglia. GUSB-deficient microglia failed to acquire homeostatic identity. Single-cell RNA sequencing identified them as a distinct population enriched for a core transcriptional program shared by immature and disease-associated microglia (DAM). Together, these findings identify lysosomal GAG catabolism as a pathway linking ECM remodeling to the acquisition of homeostatic microglial identity during CNS development.

## INTRODUCTION

The extracellular matrix (ECM) of the central nervous system (CNS) is a dynamic and highly specialized compartment that regulates neural development, synaptic plasticity, and circuit stability throughout life^1^. The brain ECM is a complex milieu of fibrillar proteins and proteoglycans and highly enriched in glycosaminoglycans (GAGs), long, negatively charged polysaccharides composed primarily of chondroitin sulfate (CS), heparan sulfate (HS), and hyaluronic acid (HA), with an estimated GAG-to-fibrous protein ratio approaching 10:1^1–5^. Sulfated GAGs represent major ECM components of the brain, constituting up to 60% of its mass during early embryonic development and 20% in the adult^6–9^. GAGs are present both as free polysaccharides and as proteoglycans that assemble into diffuse interstitial matrices and highly organized structures such as perineuronal nets (PNNs)^10–13^, specialized ECM condensations that predominantly surround parvalbumin-positive interneurons and contribute to the stabilization of mature synaptic circuits^12,14,15^. Beyond providing structural support, GAGs actively regulate numerous aspects of CNS development, including oligodendrocyte differentiation, myelination, axon guidance, growth factor signaling, and experience-dependent plasticity^16–26^.

Considerable effort has focused on defining the molecular mechanisms controlling GAG synthesis and extracellular matrix assembly. In contrast, remarkably little is known about how GAGs are removed once deposited into the extracellular environment. ECM remodeling occurs through distinct extracellular and intracellular degradative pathways. Matrix metalloproteinases (MMPs), ADAMTS proteases, and related enzymes remodel extracellular protein components of the matrix^25,27–30^, whereas degradation of CS, HS, and HA occurs predominantly following cellular uptake and lysosomal processing, with a few enzymes (HPSE, HYAL2) that are capable of extracellular GAG degradation^31^. Although the lysosomal enzymatic pathway responsible for GAG degradation has been comprehensively defined biochemically, the cellular mechanisms governing GAG uptake, intracellular degradation, and recycling within the CNS remain largely unknown. It is likewise unknown whether constitutive GAG catabolism is broadly distributed among neural cell types or concentrated within specialized cells that function as metabolic hubs for ECM turnover.

Microglia are the resident macrophages of the CNS, where they continuously survey the neural environment and internalize apoptotic cells, synapses, myelin debris, protein aggregates, and other extracellular substrates through highly active endo-lysosomal pathways^32–34^. In addition to these canonical homeostatic functions, increasing evidence indicates that microglia contribute to remodeling the GAG-rich ECM, including turnover of PNNs through both phagocytic clearance and secretion of MMPs^32,35^. Reciprocally, extracellular proteoglycans and GAGs regulate microglial activation and inflammatory signaling, suggesting a bidirectional relationship between ECM composition and microglial state^36–38^. Together, these observations raise the possibility that microglia serve as principal cellular mediators of GAG catabolism in the healthy CNS.

The importance of lysosomal GAG degradation is illustrated by mucopolysaccharidoses (MPS), a family of inherited lysosomal storage disorders caused by deficiencies in enzymes required for the degradation of GAGs^39–41^. Progressive accumulation of undegraded substrates results in severe multisystem disease accompanied by profound neurological dysfunction, including developmental delay, cognitive impairment, seizures, and premature mortality^41–45^. Collectively, these disorders demonstrate that continuous lysosomal GAG degradation is essential for CNS development and function. However, whether specific CNS cell populations are disproportionately responsible for constitutive GAG turnover—and whether selective dysfunction of these cells drives downstream neuropathology—has remained unresolved.

Mucopolysaccharidosis VII (MPS VII; Sly syndrome) results from loss-of-function mutations in β-glucuronidase (GUSB), a lysosomal glycosidase that catalyzes the terminal degradation step shared by all the major CNS GAGs, CS, HS, and HA^46–49^. Both patients with MPS VII and the well-characterized *Gusb^mps^* mouse model exhibit progressive lysosomal storage and neurological dysfunction^50–52^, providing a powerful genetic system in which constitutive GAG catabolism is selectively disrupted.

Here, we identify microglia as the principal cellular site of intracellular GAG accumulation following GUSB loss, revealing these cells as a dominant catabolic hub for constitutive extracellular matrix turnover *in vivo*. Using liquid chromatography–mass spectrometry (LC–MS), we identify that the accumulated substrate consists predominantly of chondroitin sulfate. GAG accumulation occurs within enlarged CD68+/LAMP1+ phagolysosome compartments and drives a robust, cell-intrinsic transition toward a non-homeostatic transcriptional state, indicating that GAG accumulation functions as an active instructive signal rather than passive storage material. Together, our findings identify constitutive lysosomal GAG catabolism as a previously unrecognized regulator of microglial identity and establish microglia as central mediators of ECM homeostasis in the developing brain.

## RESULTS

### Ablation of β-glucuronidase results in intracellular GAG aggregates in microglia

The lysosomal enzyme β-glucuronidase (GUSB) is an essential member of enzymatic pathways catabolizing much of the CNS GAGs, including chondroitin sulfate (CS), heparan sulfate (HS), and hyaluronic acid (HA) (Fig. S1A). To examine the consequences of GUSB loss on GAG catabolism, we used the B6.C-*H2-K^bm^*^1^/ByBir-*Gusb^mps^*/J mouse strain, which carries a null allele (*Gusb^mps^*) and serves as an established model of mucopolysaccharidosis type VII (MPS VII). Because CS-GAGs constitute ∼80% of total CNS GAGs^53^, we assessed their distribution using Wisteria floribunda agglutinin (WFA), which prominently stains for CS-GAG components of the PNN, the major ECM in the CNS. Homozygous mutants (*Gusb^mps/mps^*) lacked detectable β-glucuronidase activity (Fig. S1B), consistent with prior reports^50^ and adult *Gusb^mps/mps^* brain tissue showed pronounced accumulation of cellular WFA-positive GAG aggregates accompanied by a marked loss of extracellular WFA-labeled perineuronal nets (PNNs), the predominant ECM structures enriched around parvalbumin (PV) interneurons (Figs. 1A-C, S1C-D). We next examined the developmental onset of these aggregates. Intracellular WFA-positive aggregates were already evident in *Gusb^mps/mps^* cortex at P14, and persisted through adulthood, assessed at 4 and 9 months of age (Figs. 1A–B and S1C). Aggregate density was highest at P14 (859 ± 12.7 aggregates/mm²) and decreased with age, to 634 ± 50.0 aggregates/mm² at 4 months and 363 ± 16.0 aggregates/mm² at 9 months (Fig. 1D). Heterozygous mice (*Gusb^+/mps^*), which retain ∼50% of β-glucuronidase activity, showed no detectable intracellular WFA aggregates (Fig. S1D), indicating that complete, rather than partial, loss of enzyme activity is required to produce this phenotype. Together, these findings show that loss of lysosomal GAG catabolism leads to early-onset, persistent intracellular GAG accumulation and disrupted extracellular ECM organization in the CNS, underscoring a critical role for GUSB in CNS GAG homeostasis.

**Figure 1.**
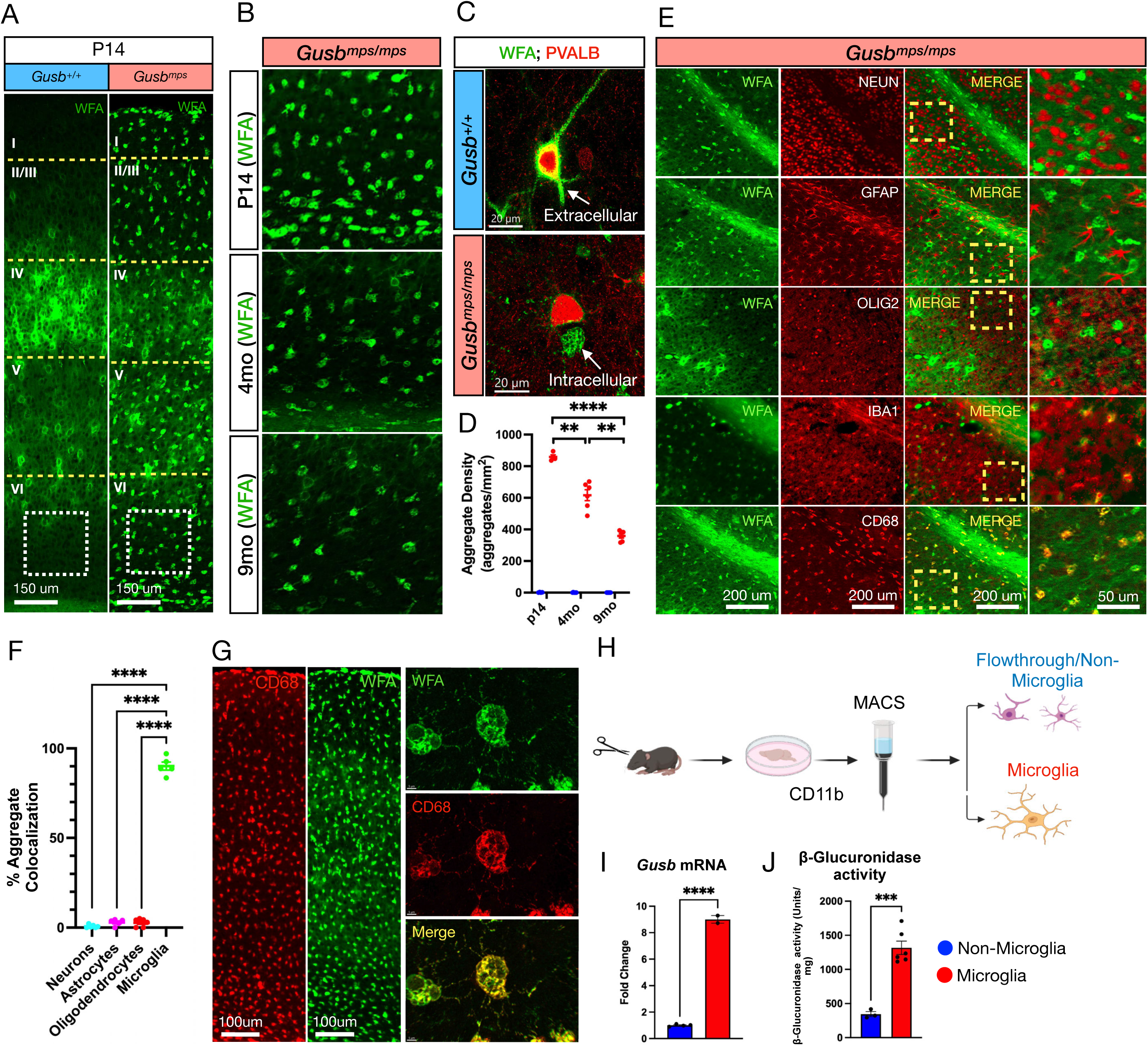
Ablation of β-glucuronidase results in intracellular glycosaminoglycan aggregates in microglia. (A) Representative WFA immunostaining (green) of the primary somatosensory cortex (SSp) from P14 *Gusb^+/+^*and *Gusb^mps/mps^* mice showing loss of perineuronal nets (PNNs) and accumulation of intracellular WFA-positive glycosaminoglycan (GAG) aggregates in mutant mice. (B) Representative images of layer VI of the SSp from *Gusb^mps/mps^* mice at P14, 4 months, and 9 months demonstrating the persistent presence of intracellular WFA-positive (green) GAG aggregates throughout aging. (C) Representative confocal images of layer IV of the SSp stained for WFA (green) and parvalbumin (PV; red), showing loss of extracellular PNNs surrounding PV+ interneurons together with intracellular WFA-positive GAG aggregates in *Gusb^mps/mps^* mice. (D) Quantification of cortical WFA-positive aggregate density (y-axis) across aging, demonstrating persistent aggregate accumulation with a progressive decline in density from P14 to 9 months (one-way ANOVA, *F*(2,10) = 63.29, *P* < 0.0001; Dunnett’s multiple-comparisons test with Tukey correction; N = 4). (E–F) Representative immunostaining (E) and quantification (F) of WFA colocalization with neuronal (NEUN), astrocytic (GFAP), oligodendroglial (OLIG2), microglial (IBA1), and lysosomal macrophage (CD68) markers in layer VI of the SSp. WFA-positive aggregates localize almost exclusively to microglia and their lysosomes (one-way ANOVA, *F*(3,18) = 1254, *P* < 0.0001; Dunnett’s multiple-comparisons test with Tukey correction; N = 3). (G) Representative CD68 (red) and WFA (green) immunostaining of the primary somatosensory cortex (SSp) at P14 at low resolution (left) demonstrating the phenotype across all layers of the SSp, and with high-resolution confocal imaging (right) confirming intracellular localization of WFA-positive GAG aggregates within microglia. (H) Schematic illustrating MACS-based isolation of neonatal cortical microglia from P14 *Gusb^+/+^* mice generated using BioRender (https://BioRender.com). (I–J) Quantification of *Gusb* mRNA expression by qRT-PCR (y-axis; fold change) (I) and β-glucuronidase enzymatic activity (y-axis; β-glucuronidase activity/mg) (J) in MACS-isolated cortical microglia and non-microglial cells from *Gusb^+/+^*mice, demonstrating enrichment of both *Gusb* expression (Student’s *t*-test, *t*(4) = 42.43, *P* < 0.0001; N = 2) and β-glucuronidase activity (Student’s *t*-test, *t*(7) = 6.670, *P* = 0.0003; N = 3) in microglia.

To determine the cellular identity of WFA-positive aggregates, we co-stained *Gusb^mps/mps^* CNS tissue with WFA and markers for neurons (NEUN), oligodendrocytes (OLIG2), astrocytes (GFAP), and microglia (IBA1 and CD68). Over 90% of GAG aggregate-containing cells co-localized with IBA1+ microglia, whereas other glial and neuronal populations, including layer IV PNN-bearing neurons, showed little intracellular GAG accumulation (Figs. 1C, E–G, and S1E). All aggregate-containing cells co-localized with intensely stained CD68, a marker of macrophage/microglial lysosomes (Fig. 1G). Thus, loss of GUSB-mediated GAG catabolism in the CNS results in abnormal intracellular GAG accumulation primarily within microglia.

These findings prompted us to assess β-glucuronidase levels in microglia relative to other CNS cell types. We measured *Gusb* mRNA expression and β-glucuronidase activity in microglia isolated from control cortical tissue by Magnetic-Activated Cell Sorting (MACS) using CD11b-conjugated beads, expressing results as fold change relative to non-microglial cells (flow-through fraction) (schematic, Fig. 1H). Microglia showed ∼8-fold enrichment of *Gusb* mRNA (Fig. 1I) and ∼3.5-fold higher β-glucuronidase activity (Fig. 1J) relative to flow-through cells, consistent with the preferential accumulation of GAG aggregates in microglia upon loss of GUSB. These results align with *Gusb* expression trends in the published Brain RNA-Seq database^54^ and the Allen Brain Cell (ABC) Atlas^55^ (Figs. S1F,H-I). Although *Gusb* is variably expressed across cell types and not exclusive to any one neural lineage, microglia show the highest overall expression (Fig. S1H). This trend was true for the 11 genes (Fig. S1G) coding for GAG degradation enzymes responsible for degrading CS, HS, and HA GAGs in the CNS. The overall expression (FPKM values) of the GAG degradation pathway was the highest in microglia (Fig. S1G).

### Lysosomal abnormalities in β-glucuronidase ablated microglia

Because GUSB is a lysosomal hydrolase, and the WFA+ accumulations appear to take up a large portion of the volume of the cells they reside in, we assessed for changes in microglial lysosomes using CD68 (also known as LAMP4), a marker of mature lysosomes and phagolysosomes in microglia, in *Gusb^mps/mps^*CNS. High-resolution imaging and 3D reconstruction of CD68+ lysosomes using Imaris revealed markedly enlarged lysosomes in *Gusb^mps/mps^* mice relative to controls (>10-fold increase in average volume; control = 0.3167 ± 0.0376 μm³, *Gusb^mps/mps^*= 5.058 ± 0.7649 μm³, p = 0.0027; Figs. 2A–B) and increased overall density of CD68 lysosomes (>3 fold higher; p<0.0001; Figs. 2A,C). Abnormal, large CD68+ lysosomes were observed exclusively in *Gusb^mps/mps^* mice, and this enlargement was a consistent feature present in both CC and cortex across all ages investigated (P14, 4 months, and 9 months) in *Gusb^mps/mps^* CNS (Fig. 2D-E and S2A).

**Figure 2.**
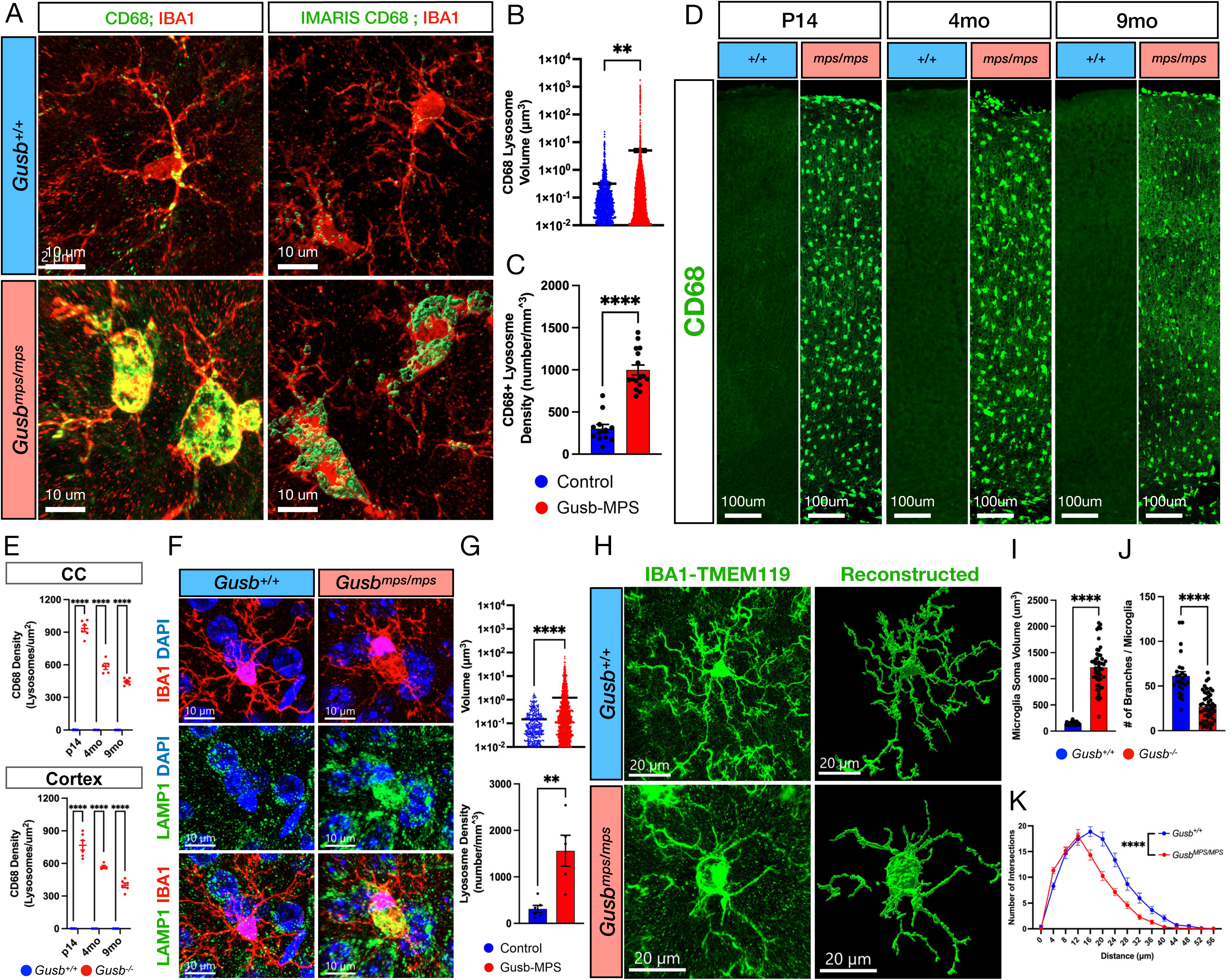
β-Glucuronidase deficiency induces lysosomal enlargement and a phagocytic microglial phenotype. (A-C) Representative images and three-dimensional Imaris reconstructions of CD68+ lysosomes (green) and IBA1+ microglia (red) from layer IV of the SSp in *Gusb^+/+^* and *Gusb^mps/mps^* mice (A), with quantification of (B) average lysosomal volume and (C) CD68+ lysosome density. *Gusb^mps/mps^* microglia exhibit a >10-fold increase in CD68+ lysosomal volume and a >3 fold increase in CD68+ lysosomal density (Student’s *t*-test: lysosomal volume, *t*(6329) = 3.001, *P* = 0.0027; lysosomal density, *t*(26) = 8.653, *P* < 0.0001; N = 3). (D-E) Representative CD68 immunostaining of the SSp from *Gusb^mps/mps^*mice at P14, 4 months, and 9 months (D) with quantification (E) of abnormal lysosomes throughout postnatal development and adulthood. A persistent, decreasing accumulation of enlarged CD68+ lysosomes is observed across the course of aging in both CC (Multiple t-test; P14, *t*(6) = 20.72, *P* < 0.0001; 4mo, *t*(6) = 17.39, *P* < 0.0001; 9mo, *t*(6) = 24.23, *P* < 0.0001; N = 3) and cortex (Multiple t-test; P14, *t*(6) = 38.99, *P* < 0.0001; 2-4mo, *t*(6) = 15.69, *P* < 0.0001; 9mo, *t*(6) = 22.66, *P* < 0.0001; N = 3). (F–G) Representative images (F) and quantification (G) of LAMP1+ lysosomes in layer IV of the SSp. *Gusb^mps/mps^*microglia exhibit a >8-fold increase in LAMP1+ lysosomal volume and a >5 fold increase in LAMP1+ lysosomal density (Student’s *t*-test: lysosomal volume, *t*(2090) = 5.101, *P* < 0.0001; lysosomal density, *t*(9) = 3.998, *P* = 0.0031; N = 3). (H–K) Representative IBA1-TMEM119 immunostaining and Imaris-based three-dimensional reconstructions of cortical microglia from 4-month *Gusb^+/+^* and *Gusb^mps/mps^* mice (H), with quantification of (I) soma volume, (J) process branching, and (K) branch complexity by Sholl analysis. *Gusb^mps/mps^* microglia exhibit enlarged somata, reduced process branching, and diminished arbor complexity, consistent with an activated phagocytic morphology (Student’s *t*-test: soma volume, *t*(76) = 14.36, *P* < 0.0001; branching, *t*(71) = 7.388, *P* < 0.0001; two-way ANOVA for Sholl analysis, *F*(14,1185) = 152.1, *P* < 0.0001; Dunnett’s multiple-comparisons test with Tukey correction; N = 3).

To corroborate lysosomal abnormalities in microglia, we co-stained CNS sections from control and *Gusb^mps/mps^* mice for the lysosomal marker LAMP1 and IBA1, and, using 3D reconstructions, quantified the number and size of LAMP1+ lysosomes in microglia. Consistent with the CD68 data, microglial LAMP1+ lysosomal volume (>8-fold increase, control = 0.148 ± 0.012 μm^3^, *Gusb^mps/mps^* = 1.217 ± 0.105 μm^3^; p < 0.0001, t-test; Fig. 2F-G, S2B) and density (>5-fold increase, control = 311.0 ± 77.82 lysosomes/mm^3^, *Gusb^mps/mps^* = 1561.0 ± 332.9 lysosomes/mm^3^; p = 0.0031, t-test; Fig. 2F-G, S2B) was markedly increased in *Gusb^mps/mps^* CNS. Furthermore, abnormally large LAMP1+ lysosomes were strongly correlated with WFA-stained GAG aggregates (Fig. S2C). Together, our findings thus far demonstrate that GUSB loss results in abnormally enlarged microglial lysosomes resulting from GAG accumulation.

Because enlarged CD68+ lysosomes are commonly associated with phagocytic, activated microglial states, we hypothesized that GUSB-deficient microglia would similarly display features of activation. Using 3D reconstructions of IBA1-TMEM119+ microglia, we found that *Gusb^mps/mps^* microglia exhibited pronounced morphological changes consistent with activation, including an amoeboid appearance (Fig. 2H), increased soma volume (>8-fold; p < 0.0001; Fig. 2I), reduced branch number (50% reduction; p < 0.0001; Fig. 2J), and decreased morphological complexity by Sholl analysis (p < 0.0001; Fig. 2K). These changes were observed broadly throughout the cortex, indicating a widespread shift toward an activated microglial state.

### G*usb*-null microglia recapitulate a phagocytic state seen in *Gusb^mps^* mice

To confirm that the phenotypes observed in the *Gusb^mps^*mouse line (B6.C-*H2-K^bm^*^1^/ByBir-*Gusb^mps^*/J), which is a major histocompatibility congenic strain, were not influenced by the congenic genetic background, we generated a new mouse line carrying a floxed *Gusb* allele using CRISPR/Cas9 (schematic, Fig. 3A) and validated the *Gusb^FLX^* allele in F2 *Gusb^FLX/FLX^* mice by genotyping (Fig. S3A) and sequencing. We crossed *Gusb^FLX/FLX^* mice with E2a-Cre (germline Cre deletor; JAX:003724) to generate heterozygous (*Gusb^+/-^*; “GUSB-HET”) and global null (*Gusb^-/-^*; “GUSB-KO” or “*Gusb* null”) mutants. *Gusb* mRNA expression and β-glucuronidase activity in CNS were reduced by ∼50% in GUSB-HET mice and by >99% in GUSB-KO mice relative to controls (Figs. 3B–C, S3B). A comparable loss of β-glucuronidase activity was confirmed in GUSB-HET and GUSB-KO in liver tissue (Fig. S3B). To confirm that residual activity in GUSB-KO tissue was non-detectable rather than below assay sensitivity, we repeated the enzymatic assay using a saturating amount of brain lysate (Fig. S3C). Thus, we validated total loss of β-glucuronidase activity in GUSB-KO (*Gusb^-/-^*) mutants generated using *Gusb^FLX^* allele with E2a-Cre. Fewer *Gusb^-/-^*mutant pups were born than expected according to Mendelian inheritance with approximately 30% of *Gusb^-/-^*pups failing to survive to birth (Fig. S3D). Next, we compared our GUSB-KO (*Gusb^-/-^*) mutants with the phenotype exhibited by *Gusb^mps^* mice. *Gusb^-/-^*mice recapitulated several characteristics previously reported in *Gusb^mps/mps^* ^52^ and *Gusb^mps2J^* ^56^ models including reduced body weight and length (Fig. 3D-F) and a stunted facial phenotype.

**Figure 3.**
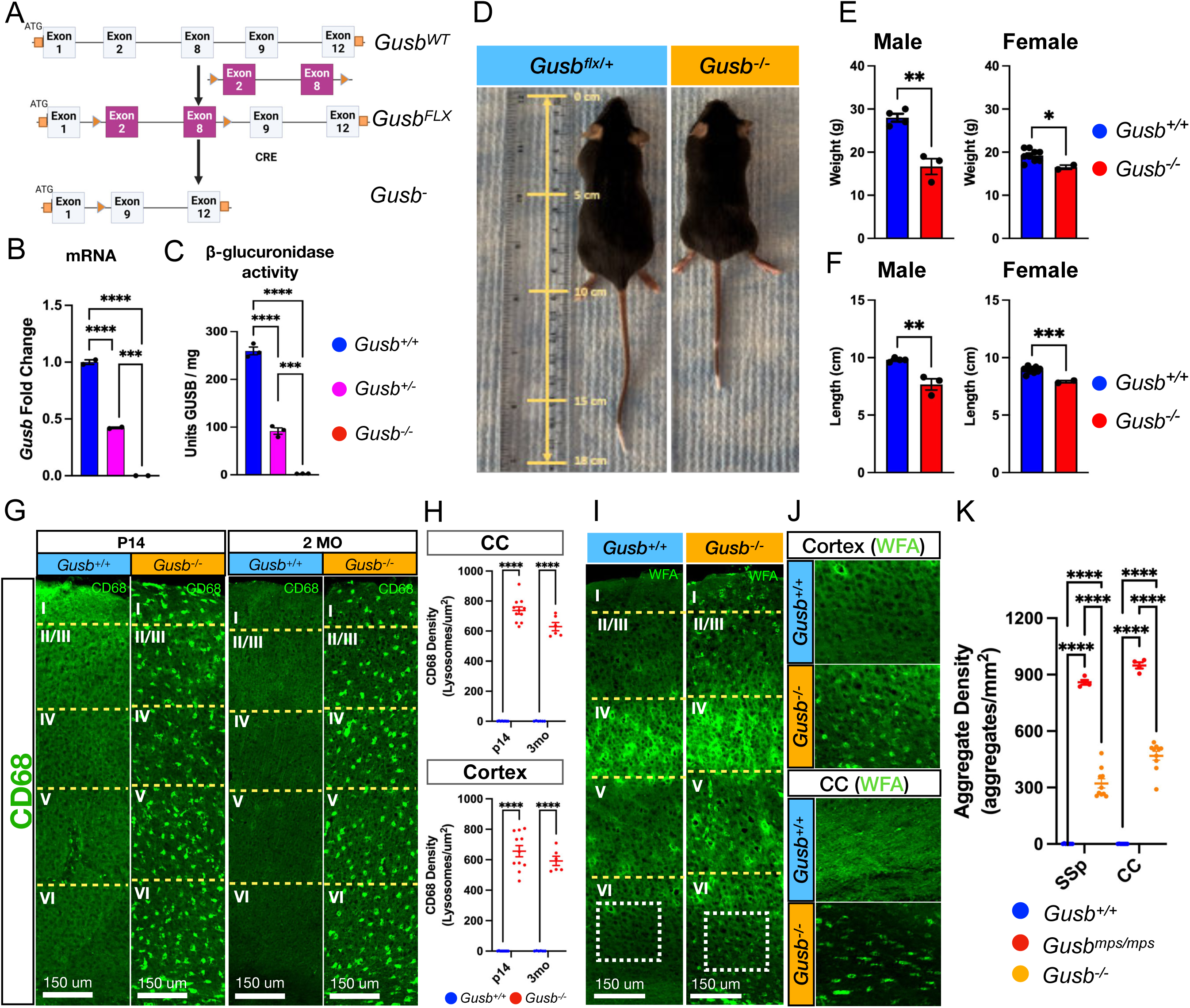
A constitutive *Gusb* knockout phenocopies the neuropathological features of the *Gusb^mps/^*^mps^ mouse. (A) Schematic illustrating generation of the *Gusb* floxed allele and constitutive *Gusb* knockout mouse line. (B–C) Quantification of *Gusb* mRNA expression by qRT-PCR (B) and β-glucuronidase enzymatic activity (C) in P14 CNS tissue from *Gusb^+/+^*, *Gusb^+/-^*and *Gusb^-/-^* mice. *Gusb^-/-^* mice exhibit complete loss of *Gusb* expression and enzyme activity relative to littermate controls (one-way ANOVA: mRNA, *F*(2,3) = ∞, *P* < 0.0001; enzyme activity, *F*(2,6) = 461.5, *P* < 0.0001; Dunnett’s multiple-comparisons test with Tukey correction; N = 2). (D–F) Representative images (D) and quantification of (E) body weight and (F) nose-to-tail length in 3-month-old *Gusb^+/+^* and *Gusb^-/-^* mice. Male and female *Gusb^-/-^* mice exhibit significantly reduced body weight (males: Student’s *t*-test, *t*(5) = 5.992, *P* = 0.0019, N = 3–4; females: *t*(9) = 2.607, *P* = 0.0284, N = 2–9) and body length (males: *t*(5) = 5.055, *P* = 0.0039, N = 3–4; females: *t*(9) = 4.973, *P* = 0.0008, N = 2–9). (G-H) Representative CD68 immunostaining (G) and corresponding quantification (H) in P14 and 2-month-old *Gusb^+/+^*and *Gusb^-/-^* mouse CC (Multiple t-test; P14, *t*(6) = 17.57, *P* < 0.0001; 2mo, *t*(10) = 18.66, *P* < 0.0001; N = 3) and SSp cortex (Multiple t-test; P14, *t*(18) = 15.98, *P* < 0.0001; 2mo, *t*(10) = 19.45, *P* < 0.0001; N = 3). (I-K) Representative WFA immunostaining (I-J) and corresponding quantification (K) of the SSp (I) from 3-month-old *Gusb^+/+^*and *Gusb^-/-^* mice, with (J) magnified views of the SSp (one-way ANOVA: *F*(2,19) = 334.0, *P* < 0.0001; Control vs MPS, P < 0.0001; Control vs KO, P < 0.0001; MPS vs KO, P < 0.0001, Tukey multiple comparisons test; N = 2–4) and CC (one-way ANOVA: *F*(2,19) = 449.2, *P* < 0.0001; Control vs MPS, P < 0.0001; Control vs KO, P < 0.0001; MPS vs KO, P < 0.0001, Tukey multiple comparisons test; N = 2–4), demonstrating the presence of WFA accumulations throughout the CNS in *Gusb^-/-^* mice, though at a lower prevalence than in *Gusb^mps/mps^* mice.

We next examined the GUSB-KO CNS for intracellular GAG aggregation and abnormal microglial CD68 volume. As in *Gusb^mps^* mice, the entire microglial population in GUSB-KO cortex and white matter displayed intensely stained enlarged CD68+ lysosomes at both juvenile (P14) and young adult (2 months) CNS (Fig. 3G-H), and comparably in both sexes (Fig. S3E). GUSB-KO cortex also showed loss of extracellular PNNs accompanied by intracellular GAG aggregation within microglia (Fig. 3I–K), another core phenotype shared with *Gusb^mps^* mice, although aggregate intensity was significantly lower in GUSB-KO than in *Gusb^mps^*tissue (Figs. 3I-J and 1A–B), suggesting potential changes in GAG content in congenic background. Together, these findings confirm that the GUSB-KO line recapitulates the core phagocytic microglial phenotype associated with loss of β-glucuronidase activity.

### Selective enrichment of chondroitin sulfate and non-degraded GAGs in *Gusb* null microglia

We hypothesized that *Gusb-*null microglia would accumulate excess GAGs with a composition distinct from that of other brain cell types. To identify the GAG species that accumulate in microglia after GUSB loss *in vivo*, we profiled chondroitin sulfate (CS) and heparan sulfate (HS), the two major GAG classes in the brain. Samples were MACS-purified microglia (MG) and the corresponding non-microglial (NMG) flow-through fractions from control (*Gusb^+/+^*), GUSB-HET (*Gusb^+/-^*) and GUSB-KO (*Gusb^-/-^*) mice. Disaccharide composition was measured by hydrophilic interaction liquid chromatography–quadrupole time-of-flight mass spectrometry (HILIC-Q-TOF-MS) with glycan reductive isotope labeling (GRIL; schematic, Fig. 4A, S4A)^57^. This platform quantifies total CS and HS disaccharides and, in the same run, non-reducing end (NRE) carbohydrates which mark incompletely degraded GAG chains^57^.

**Figure 4.**
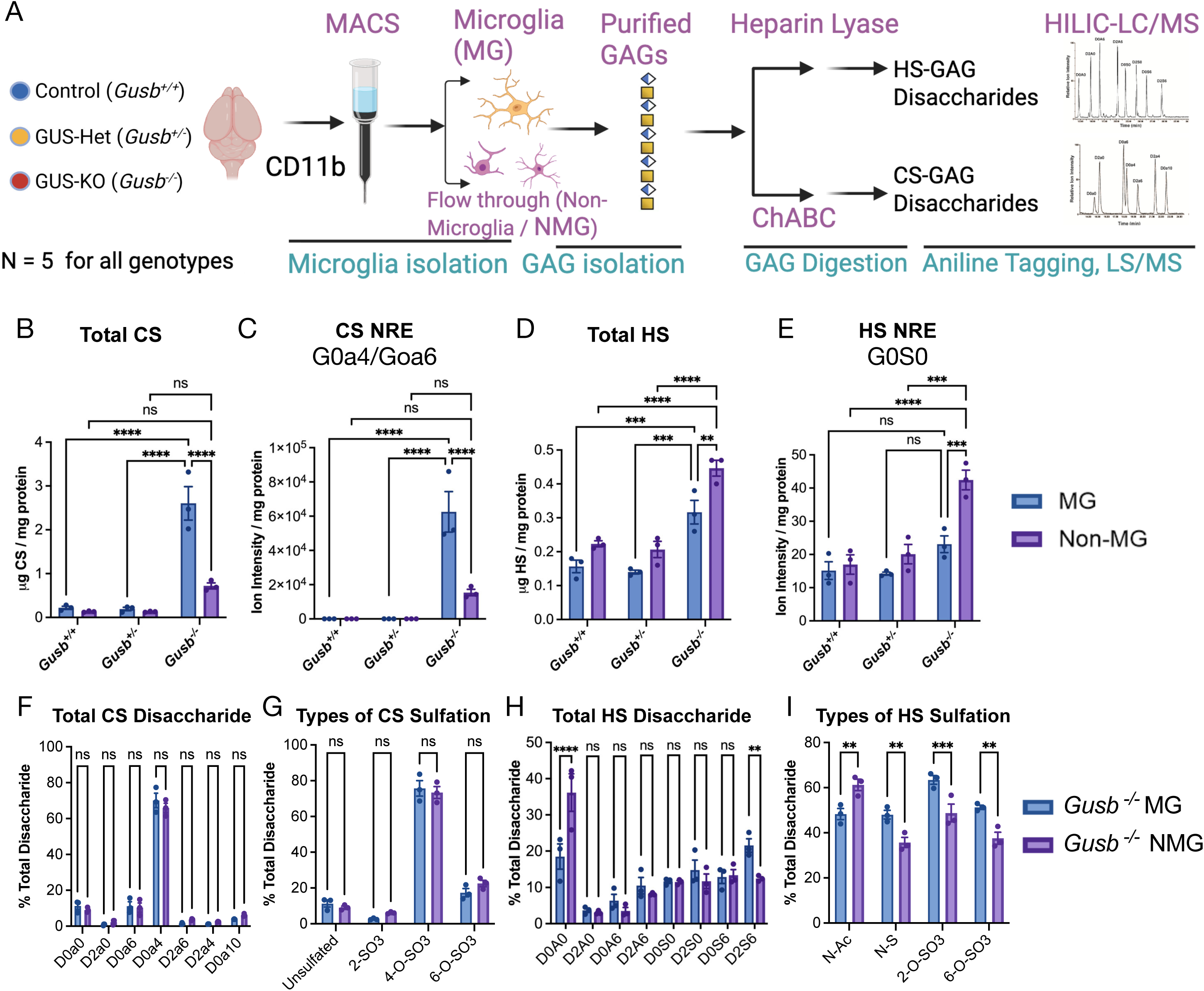
GUSB deficiency selectively promotes chondroitin sulfate accumulation and non-reducing end glycosaminoglycan species in microglia. (A) Schematic illustrating glycosaminoglycan (GAG) analysis of MACS-isolated microglia (MG) and non-microglial (NMG) cells from P14 *Gusb^+/+^*, *Gusb^+/-^*and *Gusb^-/-^* cortices generated using BioRender (https://BioRender.com). Total GAGs were isolated, digested with chondroitinase ABC or heparin lyases, and analyzed by HILIC-QTOF-MS to quantify chondroitin sulfate (CS), heparan sulfate (HS), and non-reducing end (NRE) species. (B–I) Quantification of total CS (B) and HS (D) content, corresponding CS-NRE (C) and HS-NRE (E), disaccharide composition for CS (F) and HS (H), and CS (G) and HS (I) sulfation profiles in MG and NMG isolated from *Gusb^+/+^*, *Gusb^+/-^,* and *Gusb^-/-^*cortices.

Consistent with the intracellular WFA aggregates observed in microglia, total CS-GAG was more than 10-fold higher in GUSB-KO microglia than in control or GUSB-HET microglia (two-way ANOVA, p < 0.0001; Fig. 4B). Intracellular CS-GAG was minimal in control and GUSB-HET microglia (∼0.2 µg CS-GAG/mg lysate), indicating efficient CS turnover when GUSB is present. GUSB-KO microglia also contained ∼4-fold more total CS-GAG than GUSB-KO non-microglial cells (p < 0.0001; Fig. 4B), indicating that CS accumulation is concentrated in microglia, with a smaller contribution from other cell types. Further analysis of the CS disaccharide composition (Fig. 4F) and sulfation patterns (Fig. 4G) of GUSB-KO MG and NMG suggests that CS-A or CS 4-O-sulfation is the dominant form (> 75%) of CS-GAG (Fig. 4G). As no significant changes were observed in CS-GAG disaccharide composition or sulfation patterns between MG and NMG (Fig. 4G), this suggests that microglia are not preferentially taking up and attempting to digest specific CS-GAG forms over other cell types.

HS-GAGs in GUSB-KO mice also demonstrated a distinct profile (Fig. 4D), with HS elevated ∼2-fold in GUSB-KO cells relative to control and GUSB-HET, in both microglia (KO vs. control, p = 0.006; KO vs. Het, p = 0.002) and non-microglial cells (KO vs. control and KO vs. Het, both p < 0.001). Unlike with CS-GAGs, HS-GAGs were ∼1.5-fold higher in GUSB-KO non-microglial cells than in microglia (p = 0.011; Fig. 4D). Thus, GUSB loss causes marked, microglia-selective accumulation of CS, whereas HS accumulates modestly and without microglial enrichment. Analyzing HS disaccharide composition (Fig. 4H) and sulfation pattern (Fig. 4I) in GUSB-KO MG and NMG suggested mixed changes of increased N-, 2-O- and 6-O-SO3 sulfation, and decreased N-Ac, in MG relative to NMG (Fig. 4I; Two-Way ANOVA; p < 0.01 for all).

We next asked whether the accumulated GAGs represent incompletely degraded chains. Lysosomal GAG degradation proceeds stepwise from the NRE, so loss of GUSB leaves a terminal β-linked glucuronic acid that cannot be removed. On the basis of GUSB substrate specificity, we defined the expected NRE disaccharides for each GAG class: G0a4/G0a6 (GlcA– GalNAc4S/6S) for CS and G0S0 (GlcA–GlcNS) for HS (schematic Fig. S4A). We previously validated the HS NRE G0S0 in *Gusb^mps^*cortex^57^. Here, we detected a species at m/z 553.13, consistent with G0a4/G0a6, exclusively in GUSB-KO samples, with none observed in control or GUSB-HET samples (Fig. S4B-E). This CS NRE was ∼4-fold more abundant in GUSB-KO microglia than in non-microglial cells (Fig. 4C). In contrast, G0S0 levels were similar across genotypes and cell types, except for a ∼2-fold increase in GUSB-KO non-microglial cells (vs. GUSB-KO MG, p = 0.0002; vs. Het NMG, p = 0.0001; vs. control NMG, p < 0.0001) (Fig. 4E). Together, these analyses show that GUSB deficiency causes microglia-predominant accumulation of incompletely degraded CS, identifying microglia as the principal site of impaired CS catabolism in the GUSB-deficient brain. Thus, GUSB loss drives a >10-fold, microglia-selective accumulation of CS-GAGs bearing GUSB-specific non-reducing ends, the signature of incompletely degraded chains. HS accumulates only modestly and is enriched in non-microglial cells rather than microglia, suggesting that microglia are the principal site of failed CS catabolism.

### GUSB deficient microglia are transformed into a non-homeostatic cell state with transcriptomic signature common to both naive and disease associated microglia (DAM)

To define the molecular basis of microglial abnormalities in the GUSB-deficient CNS, we performed single-cell RNA sequencing (scRNA-seq) on MACS-isolated microglia from juvenile (P21) cortex of control (*Gusb^+/+^*), GUSB-HET (*Gusb^+/-^*) and GUSB-KO (*Gusb^-/-^*) mice (schematic Fig. 5A; Methods). After quality-control filtering removed 1,430 doublets and high-mitochondrial cells (>15% mitochondrial transcripts), 27,547 of 28,977 sequenced cells were retained. Unsupervised clustering resolved six clusters (Fig. 5B, S5A) with distinct markers (Table S1). All the clusters expressed the canonical microglial markers *P2ry12, Cx3cr1, Csf1r*, and *Tgfbr1* (Fig. S5B), confirming microglial identity.

**Figure 5.**
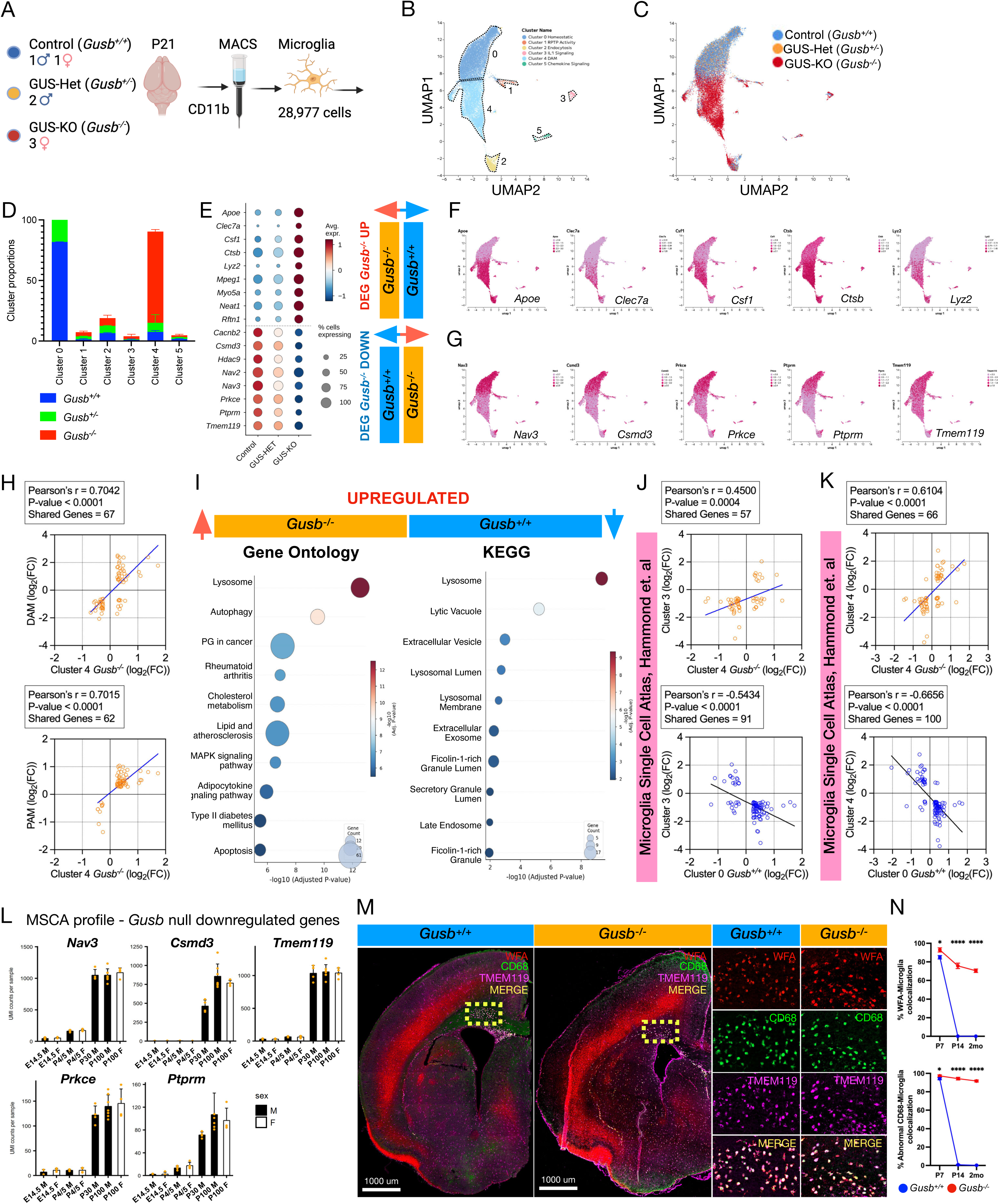
Loss of GUSB stalls the transition of developing microglia to a homeostatic cell state. (A) Schematic illustrating MACS-based isolation of cortical microglia from P21 *Gusb^+/+^*, *Gusb^+/-^*, and *Gusb^-/-^* mice for single-cell RNA sequencing generated using BioRender (https://BioRender.com). (B–C) UMAP visualization of microglial transcriptomes showing (B) six Leiden clusters, (C) the distribution of cells from each genotype across clusters. (D) Graph depicting the relative proportion of cells within each cluster. (E) Dot plot showing the expression of selected microglial marker genes across Leiden clusters. Dot size indicates the percentage of cells expressing each gene, and color intensity represents the average normalized expression level. (F–G) UMAP visualization of representative genes significantly upregulated (F) or downregulated (G) in *Gusb^-/-^*microglia relative to *Gusb^+/+^* controls. (H) Scatter plots comparing the correlation in expression, as measured by fold change compared to the unclustered data, of the matching differentially expressed genes in *Gusb^-/-^* microglia (x-axis) to those in DAM microglia (top)(r = 0.7042, P < 0.0001, Shared DEGs = 67) and PAM microglia (bottom)(r = 0.7015, P < 0.0001, Shared DEGs = 62) (y-axis). (I) Bubble plot showing KEGG and Gene Ontology pathway enrichment among genes upregulated in *Gusb^-/-^* microglia relative to *Gusb^+/+^*controls. Bubble size indicates the number of differentially expressed genes associated with each pathway, and color intensity represents enrichment significance (false discovery rate-adjusted *P* value). (J-K) Scatter plots comparing the correlation in expression, as measured by fold change compared to the unclustered data, of the matching differentially expressed genes between *Gusb^-/-^* cluster 4 (top) and *Gusb^+/+^*cluster 0 (bottom) microglia (x-axis) to Microglia Single Cell Atlas cluster 3 (J) (*Gusb^-/-^*; r = 0.4500, P = 0.0004, Shared DEGs = 57, *Gusb^+/+^*; r = - 0.5434, P < 0.0001, Shared DEGs = 91) and cluster 4 (K) (*Gusb^-/-^*; r = 0.6104, P < 0.0001, Shared DEGs = 66, *Gusb^+/+^*; r = -0.6656, P < 0.0001, Shared DEGs = 100) microglia (y-axis). (L) Bar graphs of the normal expression patterns in microglia from male and female mice across various time points for some of the top downregulated genes in *Gusb^-/-^* microglia relative to *Gusb^+/+^*controls (in order from left to right, top to bottom), genes downregulated in *Gusb^-/-^* microglia are highly expressed in adulthood. Graphs were generated using the Microglial Single Cell Database^74^. (M-N) Representative images (M) and corresponding quantifications (N) of coronal brain sections from P7 *Gusb^+/+^* and *Gusb^-/-^* mice immunostained for WFA (red), CD68 (green), and TMEM119 (magenta), with higher-magnification images of the corpus callosum demonstrating comparably similar colocalization of microglia in *Gusb^+/+^* and *Gusb^-/-^*mice with WFA+ accumulations and abnormal CD68 lysosomes (Student’s *t*-test: WFA, *t*(6) = 2.705, *P* = 0.0353, N=2; CD68, *t*(6) = 2.758, *P* = 0.0330, N=2), at P7, relative to later time points (two-way ANOVA Interactions; WFA, F(2, 18) = 203.8, P < 0.0001; CD68, F(2, 18) = 1337, P < 0.0001, N=2).

Genotype strongly determined cluster proportions (χ² test, p < 0.0001; Fig. 5C,D). Two major clusters (0 and 4) contained 88.2% of all cells. Most control and GUSB-HET microglia fell in cluster 0 (81.8% ± 0.9% and 82.0% ± 4.1% of each genotype, respectively), whereas 75.4% ± 1.0% of GUSB-KO microglia fell in cluster 4 (Fig. 5D). Cluster 0 expressed the highest levels of canonical homeostatic markers *Tmem119, P2ry12, Cx3cr1* and *Tgfbr1,* and showed highest expression of genes (*Nav3, Nav2, Csmd3, Ptprm, Prkce*) expressed only in adult homeostatic microglia (Fig. 5B,E,G, S5C and Table S1), a profile determined from the Microglial Single Cell Atlas database (MSCA) (Fig. 5L). Cluster 0 was therefore designated as the homeostatic cluster. These genes were downregulated in the GUSB-KO–dominated cluster 4, which instead upregulated genes identified in disease-associated microglia (DAM)^58^ including *Apoe, Clec7a, Csf1, Ctsb and Lyz2* (Fig. 5B,E,F, S5D and Table S2). Cluster 4 was therefore designated as the DAM cluster. Cluster 2 contained similar proportions of all three genotypes. Of the four minor clusters (3261 cells combined), clusters 1 and 3 were disproportionately GUSB-KO (46.6% and 61.3% of cells in each cluster), while cluster 5 was genotype-balanced (Fig. 5D, S5E). These minor clusters were enriched for distinct pathways: receptor protein phosphatase activity (cluster 1), endocytosis (cluster 2), IL-1 signaling (cluster 3), and chemokine signaling (cluster 5).

Further analysis of differentially expressed genes (DEGs) in GUSB-KO cells relative to control and GUSB-HET cells (FC > 1.5 and adjusted *p* < 0.02; Table S3) revealed that upregulated genes were enriched for terms related to lysosomal, lytic vacuole, and extracellular vesicle components, as well as innate immune responses, neutrophil degranulation, and cholesterol transport (Fig. 5I; Table S4). This lysosomal signature is consistent with the enlarged CD68+ and LAMP1+ volume and increased density observed in *Gusb^mps^* microglia (Fig. 2A–C, F–G). Genes downregulated in the GUSB-KO cluster were enriched for RHO GTPase signaling and focal adhesion pathways (Fig. S5F; Table S4).

### GUSB-KO signature resembles immature, early postnatal microglia

Strikingly, the profile of DEGs strongly upregulated in GUSB-KO resembled two previously described microglial cell states: disease-associated microglia (DAM)^58^, found in adult neurodegeneration models, and proliferative-region-associated microglia (PAM)^59^, found transiently in early postnatal white matter (Fig. 5H). Among the DEGs identified in our GUSB-KO, 67 overlapped with the DAM signature and 62 with the PAM signature (Fig. 5H, Table S2), and 21 were shared by both (Table S2). Thus, GUSB loss shifts juvenile microglia from a homeostatic toward a DAM/PAM-like state. The enrichment of a non-homeostatic microglial signature among GUSB-KO DEGs suggested that these microglia fail to acquire, or depart from, a mature homeostatic identity. To determine which microglial state the GUSB-KO profile most closely resembles, we correlated GUSB-KO cluster 4 DEGs with 13 developmental, aging and injury-associated microglial clusters from the MSCA database, spanning E14.5, P4/5, P30, P100, and P540 (Pearson correlation of log2FC for shared DEGs; Figs. 5J–K, S6A–B).

GUSB-KO microglia correlated most strongly with embryonic and early postnatal clusters (MSCA clusters 1–6; Figs. S6A–B). The strongest correlations were with E14.5 microglia (MSCA cluster 3: r = 0.4500, P = 0.0004, 57 shared DEGs) and P4/5 phagocytic, axon tract– associated microglia (MSCA cluster 4: r = 0.6104, p < 0.0001, 66 shared DEGs) (Fig. 5J-K), with these immature microglia clusters also being inversely correlated with control microglia cluster 0 (atlas cluster 3: r = −0.5434, P < 0.0001, 91 shared DEGs; atlas cluster 4: r = −0.6656, p < 0.0001, 100 shared DEGs) (Fig. 5J-K). Interestingly, the GUSB-KO DEGs shared with the P4/5 axon tract–associated cluster were also enriched for lysosomal pathways (Fig. S6C-D) and included *Apoe*, *Cd68*, *Lamp1*, *Csf1*, and *Lyz2*, genes also shared by DAM and PAM, suggesting that these microglial cell states share a common signature of increased phagocytic activity.

Yet another consistent theme to emerge from the comparisons of the GUSB-KO microglia with the MSCA database is that the most strongly downregulated genes in GUSB-KO (*Nav3*, *Csmd3*, *Tmem119*, *Prkce*, and *Ptprm*) are highly expressed in P30 and P100 homeostatic microglia (Fig. 5L), while the most strongly upregulated genes (*Apoe, Arhgap24, Csf1* and *Myo5a*) were highly expressed in E14.5 and P4/5 microglia (Fig S6E). Together, this data suggests that GUSB-deficient microglia resemble an immature, phagocytic state similar to P4/5 microglia.

### Early postnatal wild-type microglia transiently accumulate GAGs

The resemblance to early postnatal microglia led us to ask whether GAG uptake and lysosomal processing are normal features of microglia at this stage. We therefore examined WFA+ accumulations and enlarged CD68+ lysosomes in P7 white matter. Wild-type microglia already showed extensive WFA+ accumulations and prominent CD68+ lysosomal aggregates, closely resembling mutant microglia. Microglial colocalization with both features was high in both genotypes and only modestly greater in mutants (WFA+: *Gusb^+/+^* 84.98% vs. *Gusb^-/-^*92.91%, p = 0.0353; enlarged CD68+ lysosomes: 94.18% vs. 98.05%, p = 0.0330; Fig. 5M–N). By P14 the genotypes diverged: as these features declined in wild-type microglia but persisted in GUSB-deficient microglia (two-way ANOVA, genotype × age interaction, p < 0.0001 for both measures; Fig. 5M–N). These findings indicate that microglia physiologically take up and process GAGs during early postnatal development. They further suggest that GUSB-dependent GAG catabolism is required for microglia to exit this phagocytic state and adopt a homeostatic identity.

### Persistent DAM and phagocytic marker expression following loss of GUSB activity

We next examined the temporal expression of microglia-enriched genes identified by scRNA-seq as among the most significantly altered in GUSB-KO mice, including markers of DAM and PAM (Clec*7a*), phagocytic lysosomes (*Cd68, Mpeg1*), and chemokine signaling (*Ccl3, Ccl4*). Expression was assessed in cortical tissue from GUSB-KO and control mice at P14, P21, and 2 months. All genes examined were significantly upregulated at every age tested, though the magnitude of induction varied by gene (Fig. 6A), indicating persistent elevation of DAM- and phagocytosis-associated transcriptional programs from early postnatal development into adulthood. A nearly identical pattern was observed in white matter (Fig. 6B), indicating that this transcriptional signature is distributed broadly across both gray and white matter. These findings are consistent with the widespread CD68+ enlarged microglia observed histologically (Fig. 2A-E and 3G–H).

**Figure 6.**
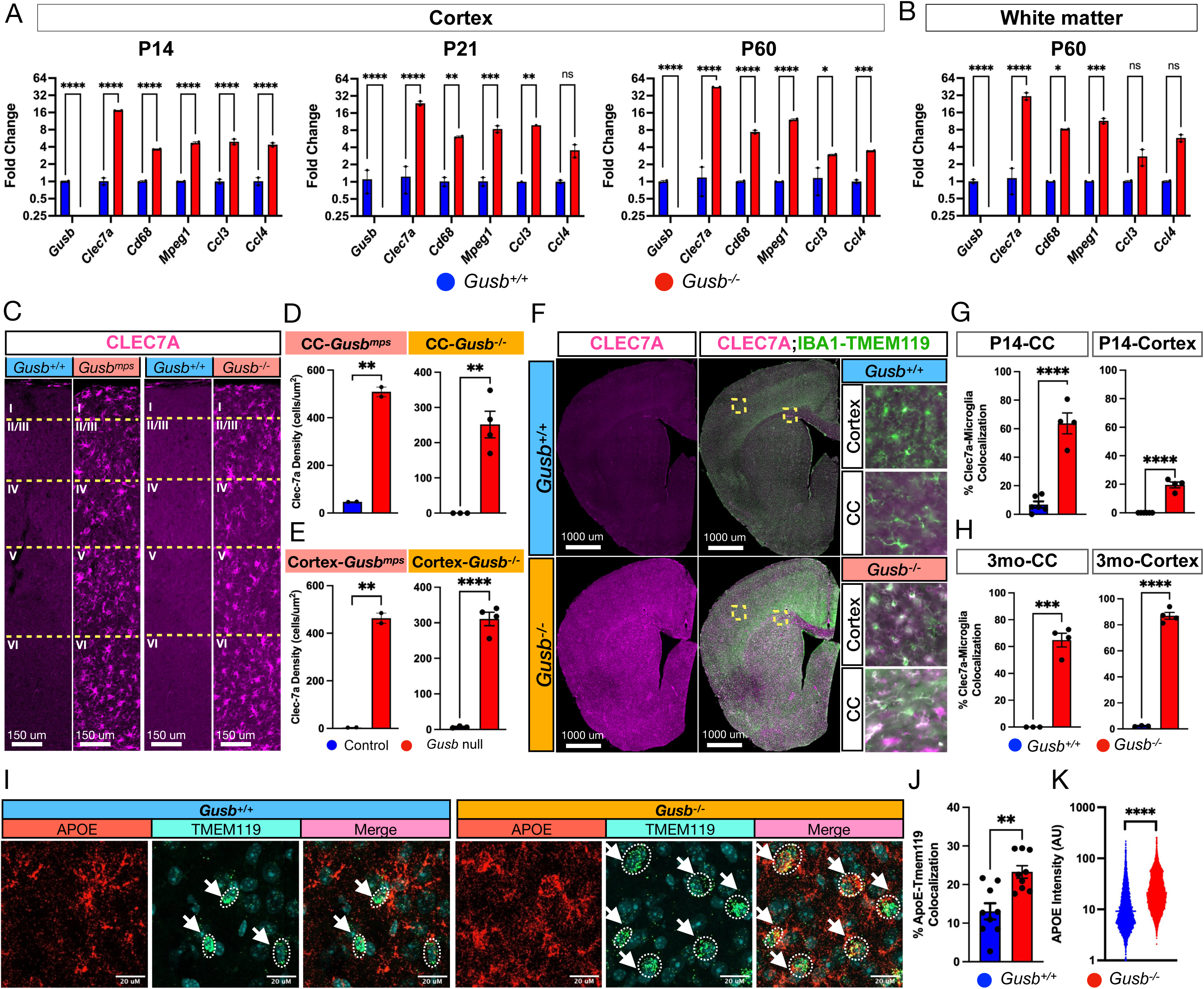
Loss of GUSB induces persistent disease-associated microglial marker expression throughout postnatal development. (A–B) qRT-PCR analysis of disease-associated microglial (DAM) marker expression in *Gusb^-/-^* and *Gusb^+/+^*mice. (A) Relative gene expression in cortical tissue at P14, P21, and 2 months, and (B) white matter at 2 months. *Gusb^-/-^* mice exhibit sustained upregulation of DAM-associated genes throughout postnatal development (one-way ANOVA: P14, *F*(5,12) = 429.9, *P* < 0.0001; P21, *F*(5,12) = 51.01, *P* < 0.0001; 2 months cortex, *F*(5,12) = 1459, *P* < 0.0001; white matter, *F*(5,12) = 33.98, *P* < 0.0001; Dunnett’s multiple-comparisons test with Tukey correction; N = 1–2). (C-E) (C) Representative images of CLEC7A immunostaining from layers 1-6 of the SSp from Control, *Gusb^mps/mps^,* and *Gusb^-/-^*mice at 3 months of age. (D-E) Quantifications of CLEC7A+ microglial density from *Gusb^mps/mps^* mice (left) and *Gusb^-/-^*mice (right) in CC (D) and SSp (E). In both *Gusb^mps/mps^* and *Gusb^-/-^* mice, we observe a large increase in CLEC7A+ microglial density (Student’s *t*-test: *Gusb^mps/mps^* corpus callosum, *t*(2) = 22.93, *P* = 0.0019; *Gusb^-/-^* corpus callosum, *t*(5) = 5.603, *P* = 0.0025; *Gusb^mps/mps^* cortex, *t*(2) = 21.37, *P* = 0.0022; *Gusb^-/-^* cortex, *t*(5) = 13.49, *P* < 0.0001; N = 2). (F) Representative coronal brain sections co-immunostained for CLEC7A (magenta) and IBA1-TMEM119 (green), with higher-magnification images of the cortex and corpus callosum demonstrating selective expression of CLEC7A in microglia from *Gusb^-/-^* mice. (G-H) Quantification of CLEC7A colocalization with IBA1+ microglia in the corpus callosum and cortex at (G) P14 and (H) 3 months. CLEC7A expression is significantly increased in microglia from *Gusb^-/-^* mice at both ages (Student’s *t*-test: P14 corpus callosum, *t*(8) = 8.797, *P* < 0.0001; P14 cortex, *t*(8) = 12.00, *P* < 0.0001; 3-month corpus callosum, *t*(5) = 10.74, *P* = 0.0001; 3-month cortex, *t*(5) = 27.05, *P* < 0.0001; N = 4). (I-K) Representative RNAscope images (I) and quantification of (J) APOE/TMEM119 colocalization and (K) APOE signal intensity in cortical microglia. *Gusb^-/-^* mice exhibit increased APOE expression within TMEM119+ microglia relative to *Gusb^+/+^*controls (Student’s *t*-test: APOE/TMEM119 colocalization, *t*(16) = 3.848, *P* = 0.0014; APOE intensity, *t*(15385) = 40.24, *P* < 0.0001; N = 3).

We next immunostained for CLEC7A (Dectin-1), a pattern-recognition receptor and hallmark marker of DAM and PAM. In adult GUSB-KO mice, CLEC7a was robustly expressed by IBA1+ microglia in both cortex and white matter but was essentially absent in control littermates (Fig. 6C–H, S7A). Like CD68+ lysosomal enlargement, CLEC7A induction is therefore a consistent consequence of GUSB deficiency. The CLEC7A+ phenotype emerged at different times in the two regions. In white matter, ∼70% of IBA1-TMEM119+ microglia were CLEC7A+ at both P14 and 3 months. In cortex, this proportion rose from ∼25% at P14 to ∼90% at 3 months (Fig. 6F– H, S7B). Additionally, as with prior observations of WFA+ accumulations and prominent CD68+ lysosomes in P7 white matter in *Gusb^+/+^* mice, we identify extensive expression of CLEC7A within microglia at P7 as well, with no significant difference observed in the density of CLEC7A+ microglia between *Gusb^+/+^* and *Gusb^-/-^*mouse white matter (CLEC7A+: *Gusb^+/+^* 194.8 cells/mm^2^ vs. *Gusb^-/-^*285.4 cells/mm^2^, p = 0.0686; Fig. S7C-D). Thus, GUSB deficiency induces a sustained non-homeostatic DAM and PAM-like phenotype that begins in early postnatal white matter and later extends to the cortex.

*Apoe*, another hallmark DAM gene identified by scRNA-seq, is expressed mainly by astrocytes rather than microglia under physiological conditions. RNAscope for *Apoe* and *Tmem119* in 2-month-old cortex showed that GUSB-KO microglia had higher APOE intensity (p < 0.001) and greater APOE/TMEM119 colocalization (p < 0.01; t-tests; Fig. 6I-K) than controls, throughout the cortex, paralleling the CD68 and CLEC7A phenotypes. Together, these findings show that GUSB loss drives a persistent shift toward a DAM-like state, marked by sustained induction of canonical DAM genes including *Clec7a* and *Apoe*.

### GUSB ablation and CSPGs trigger abnormal phagocytosis in purified microglia

Testing whether GUSB acts cell-autonomously requires microglia-specific deletion. However, lysosomal enzymes are exported and taken up by neighboring cells via the mannose-6-phosphate receptor, a mechanism known as cross-correction, which is particularly well established for GUSB^60,61^. Cross-correction underlies the efficacy of bone marrow transplantation, gene therapy and enzyme replacement therapy in MPS VII mice and patients^60,62–64^. Because secreted GUSB from neighboring cells could therefore mask a microglia-specific deletion, we used purified microglial cultures from GUSB-KO mice to test whether GUSB loss cell-autonomously increases phagocytosis, a function suggested by the lysosomal and phagocytic signatures of GUSB-KO microglia *in vivo*.

We thus assessed phagocytic activity from primary microglia isolated from neonatal (P6) cortices of control (*Gusb^+/+^*) and GUSB-KO (*Gusb^-/-^)* mice with and without LPS (100 ng/mL) and/or CSPG (100 µg/mL) treatments (experimental schematic, Fig. 7A). We confirmed the purity of our microglial isolation by optimizing our protocol using cortical tissue derived from CX_3_CR1-GFP knock-in mouse strain (B6.129P2(Cg)-*Cx3cr1^tm1Litt^*/J). Microglia derived from CX_3_CR1-GFP knock-in cortex were ∼ 95% positive for CX_3_CR1-GFP, a marker for microglia and 99% IBA1 positive (Fig. S8A-B), suggesting high purity of microglia using this protocol (Materials and Methods). Additionally, we measured β-glucuronidase activity and confirmed the total ablation of enzymatic activity in glia derived from GUSB-KO cortex in our cultures (Fig. S8C).

**Figure 7.**
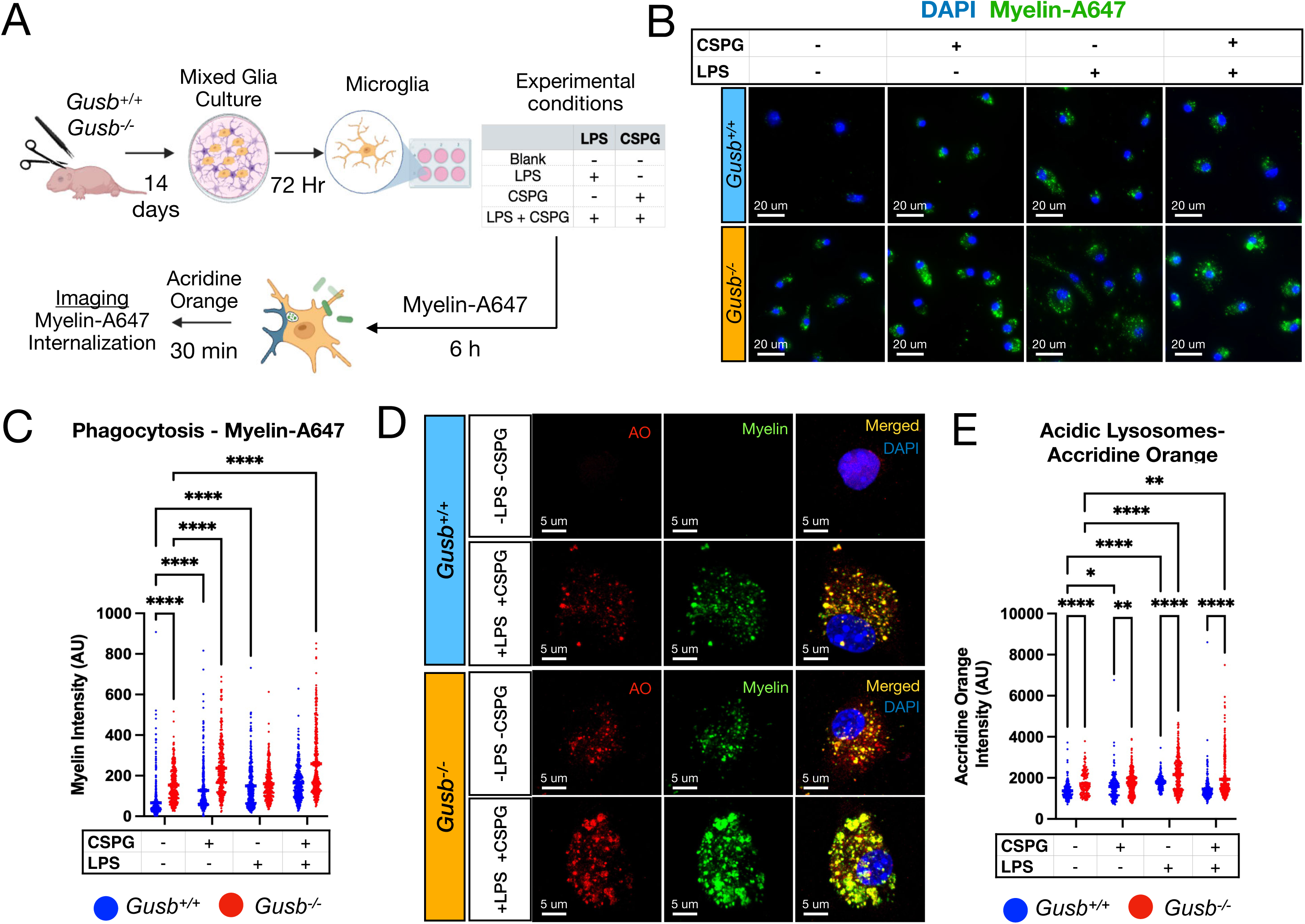
Loss of GUSB enhances microglial phagocytic activity and is further potentiated by chondroitin sulfate proteoglycans. (A) Schematic illustrating the isolation and culture of primary cortical microglia from neonatal *Gusb^-/-^* and *Gusb^+/+^*mice for *in vitro* phagocytosis assays generated using BioRender (https://BioRender.com). (B–C) Representative images (B) and quantification (C) of myelin phagocytosis by primary microglia sourced from *Gusb^+/+^*and *Gusb^-/-^* mice, treated with vehicle, lipopolysaccharide (LPS), chondroitin sulfate proteoglycans (CSPGs), or LPS plus CSPGs. *Gusb^-/-^* microglia exhibit significantly increased myelin phagocytosis under basal conditions, which is further enhanced by LPS, CSPGs, and combined LPS+CSPG treatment relative to untreated *Gusb^+/+^*controls. Statistical analysis is shown in figure S8D. (D-E) High-resolution confocal images (D) and quantification (E) of primary *Gusb^+/+^*and *Gusb^−/−^* microglia demonstrating both colocalization of internalized myelin-A647 (green) with acridine orange-labeled lysosomes (red), confirming lysosomal trafficking of phagocytosed myelin, and increased staining by Acridine Orange in microglia, demonstrating increased presence of acidic lysosomes. Statistical analysis is shown in figure S8E.

Phagocytic capacity was assessed by quantifying uptake of fluorescently labeled myelin (Myelin-A647) following a 6-hour incubation (representative images, Fig. 7B; quantification, Fig. 7C). Under basal conditions, GUSB-KO microglia exhibited a significant >3-fold increase in myelin uptake compared to untreated controls (p < 0.0001; one-way ANOVA with multiple comparisons; Fig. 7B-C). In contrast, control microglia displayed minimal baseline phagocytosis. LPS treatment robustly induced phagocytosis in control cells (∼3-fold increase relative to untreated; ****, p < 0.0001), reaching levels comparable to untreated GUSB-KO microglia (ns; Fig. 7B-C). Notably, LPS treatment produced only a modest additional increase in GUSB-KO microglia (Fig. 7B-C), indicating that GUSB deficiency phenocopies an LPS-activated state. Pairwise statistical analysis across all conditions (Fig. S8D) further supports this interpretation. Untreated control microglia differed significantly from nearly all other groups (****), whereas comparisons among LPS-treated and GUSB-KO conditions frequently did not reach significance (ns), suggesting convergence toward a common high-phagocytic phenotype. To confirm that Myelin-A647 signal represented bona fide phagocytosis, cells were stained with Acridine Orange (AO) to label acidic lysosomal compartments (Fig. 7D). Internalized myelin showed near-complete co-localization with AO-positive vesicles, indicating lysosomal trafficking. Moreover, conditions with increased myelin uptake (e.g., GUSB-KO) exhibited a corresponding increase in AO-positive vesicles (Fig. 7D-E, S8E), consistent with enhanced lysosomal activity.

Furthermore, we examined whether CSPG alone is sufficient to modulate phagocytosis. Plating microglia on CSPG-coated coverslips for 48 hours significantly increased phagocytic activity in control cells (p < 0.0001), reaching levels comparable to LPS treatment alone (ns; Fig. 7B,C). Importantly, CSPG further enhanced phagocytosis in GUSB-KO microglia (∼3-fold increase; p < 0.0001), demonstrating an additive effect of extracellular GAG exposure on an already elevated baseline. Across conditions, CSPG-treated groups frequently showed no significant difference when compared to LPS-treated groups (ns; Fig. 7B-C), reinforcing the conclusion that CSPG is a potent inducer of phagocytosis. Combined LPS and CSPG treatment resulted in only modest additional increases beyond either treatment alone in both genotypes, suggesting a ceiling effect. This plateau is further supported by the prevalence of non-significant comparisons among high-activity conditions in the statistical matrix (Fig. S8D).

Collectively, these data demonstrate that loss of GUSB is sufficient to drive elevated microglial phagocytic capacity under basal conditions. Furthermore, CSPG strongly promotes phagocytosis to an extent comparable to LPS stimulation, with a markedly amplified effect in GUSB-deficient microglia. These findings indicate that GUSB ablation and presence of excess CS-GAGs synergistically enhance microglial phagocytic activity.

## DISCUSSION

In this study, we establish that lysosomal GAG catabolism is essential for maintaining microglial homeostasis in the CNS, and that its disruption drives a cell-intrinsic transition to a DAM-like state through the accumulation of undegraded GAGs. By combining a classical MPS VII mouse model with a newly generated conditional knockout system, we provide strong genetic evidence that β-glucuronidase-mediated GAG degradation is a fundamental regulator of microglial identity. Strikingly, despite global deletion of GUSB, microglia emerged as the primary site of intracellular GAG accumulation and the principal cell type driving the neuropathological consequences of enzyme deficiency. These findings identify lysosomal GAG catabolism as a previously underappreciated checkpoint controlling microglial cell state, with implications extending beyond mucopolysaccharidosis VII to the broader biology of ECM-microglia interactions during development, aging, and neurodegeneration.

### Microglia-mediated GAG catabolism during development

A central finding of our study is the remarkable cell-type specificity of GAG accumulation. Although GUSB is ubiquitously required for GAG degradation, virtually all intracellular WFA-positive GAG aggregates localized to microglia/macrophages. LC-MS analyses further confirmed these observations by demonstrating selective accumulation of chondroitin sulfate-derived, but not heparan sulfate-derived, GAGs within microglia. This selectivity cannot be explained simply by substrate availability, since all CNS cell types reside within a GAG-rich extracellular environment. Rather, our observation that microglia exhibit substantially higher *Gusb* expression and β-glucuronidase activity than other CNS cell types suggests that they function as a major catabolic hub for ECM GAG turnover, analogous to their established roles in clearing myelin debris, synaptic material, and apoptotic cells. The appearance of intracellular GAG aggregates as early as P7 further indicates that microglia are actively engaged in GAG turnover during early postnatal development, a period of rapid ECM remodeling and circuit maturation. Strikingly, even wild-type microglia in P7 white matter contained abundant WFA+ GAG accumulations and enlarged CD68+ lysosomes, nearly indistinguishable from GUSB-KO microglia at this age. These microglia occupy the same developing white matter tracts at the same stage as PAM and P4/5 axon tract–associated microglia, whose lysosomal and phagocytic signature (including *Cd68, Lamp1, Apoe,* and *Lyz2*) closely matched that of GUSB-KO microglia. Thus, the hallmarks of GAG storage in GUSB-KO microglia are normal, transient features of early postnatal microglia that resolve by P14 in wild-type mice but persist in the absence of GUSB. These findings suggest that continual degradation of extracellular GAGs represents a constitutive housekeeping function of developing microglia. Consequently, loss of GUSB unmasks this previously unrecognized catabolic burden, resulting in a microglial lysosomal storage phenotype rather than generalized intracellular storage across CNS cell types.

### GAG-mediated microglial activation resembles both DAM and naïve postnatal microglia

The DAM program, first described in Alzheimer’s disease and other neurodegenerative disorders, is thought to reflect microglial responses to protein aggregates and neuronal loss. A closely related signature, with increased *Apoe*, *Clec7a*, and *Csf1* and reduced *Tmem119*, *P2ry12*, and *Cx3cr1*, defines proliferative-region-associated microglia (PAM) in early postnatal development. How these two states are mechanistically related remains unresolved.

Our scRNA-seq analyses show that GUSB-deficient microglia adopt a program resembling both DAM and PAM. This transition occurs by P14, before overt neurodegeneration, amyloid or tau pathology, or full circuit maturation. A DAM-like program can therefore arise independently of classical neurodegenerative triggers. Because lysosomal dysfunction is common across neurodegenerative disorders, disrupted GAG homeostasis may be a shared upstream mechanism underlying both developmental PAM and pathological DAM states. By uncoupling lysosomal dysfunction from neuronal degeneration, the GUSB-deficient mouse offers a simplified genetic model for dissecting how disease-associated microglial states are established. Whether abnormal GAG catabolism and lysosomal stress act together or independently to drive non-homeostatic states remains to be determined.

### Microglia in MPS

Our findings reframe MPS VII neuropathology as a primary microglial lysosomal disorder with secondary effects on the ECM and PNN-dependent circuits, rather than a uniform consequence of tissue-wide GAG storage. Microglia have the highest *Gusb* expression and GUSB activity among CNS cells. WFA staining and LC-MS of purified microglia show that intracellular CS storage, including non-degraded species, is concentrated in this population. Thus, microglia appear uniquely dependent on continuous lysosomal GAG degradation and serve as a major catabolic hub for ECM homeostasis.

This microglial predominance may be a general feature of MPS. Early, progressive microgliosis has been reported across MPS I, IIIA, and IIIB^65,66^, and in a recent study of *Sgsh*-deficient (MPS IIIA) mice, microglia were the most profoundly affected brain cell type by ultrastructural, transcriptomic, and epigenetic measures^67^. Our analysis of purified cell fractions provides direct biochemical evidence that undegraded GAGs accumulate preferentially in microglia.

The resulting state transition is also similar. In MPS IIIA, 83% of *Sgsh*-deficient microglia occupied a DAM/MGnD-like cluster; here, 75.4% of *Gusb*-null microglia occupied a distinct non-homeostatic cluster with markers similar to DAM, PAM and axon tract-associated microglia. Different substrates (HS in MPS IIIA, predominantly CS in MPS VII) converge on this shared program, suggesting that lysosomal failure itself, rather than a specific GAG species could be the source of this microglial state^68^.

Whether GUSB acts cell-autonomously in microglia will require microglia-specific *Gusb* ablation. However, GUSB is efficiently secreted and taken up by neighboring cells via the mannose-6-phosphate receptor. This cross-correction underlies the efficacy of bone marrow transplantation, gene therapy, and enzyme replacement therapy in MPS VII^60,62–64^. Microglia may themselves be a key source of cross-correcting enzyme: myeloid-restricted gene therapy is effective in MPS IIIA^69^, and microglia supply HEXB to neurons in Sandhoff disease^70^. Because enzyme from neighboring cells could mask a microglia-specific deletion, resolving cell autonomy may require strategies that also limit cross-correction.

### Role of CS-GAG homeostasis in regulating microglial state

Our study identifies abnormal accumulation of CS-derived GAGs within enlarged lysosomes as a defining feature of GUSB-deficient microglia transitioning to a DAM-like state. CS-GAGs are not inert storage material but bioactive ligands that instruct microglial state through innate immune receptors including TLR4 and CD44, with effects that are strikingly context- and structure-dependent. In the injured spinal cord, CSPGs signal via TLR4 to hold myeloid cells in a pro-inflammatory state and prevent their conversion to a pro-resolution phenotype^36^, yet CSPG deposition after injury also has beneficial effects on microglia/macrophage activation^71^, and CS disaccharide degradation products drive a phagocytically competent but anti-inflammatory, IFN-γ counter-regulatory phenotype distinct from classical activation^72^. Sulfation position is also a key determinant of this bidirectionality, with the 4-O-sulfated disaccharide Di-4S exerting substantially stronger modulation of TLR-dependent cytokine output than its 6-O-sulfated or non-sulfated counterparts^73^. Consistent with this, exogenous CSPG elevated phagocytic activity in wild-type microglia to a degree indistinguishable from LPS and further amplified the already elevated baseline of GUSB-KO cells, indicating that extracellular CS exposure and intracellular CS storage converge on a common phagocytic program. We therefore propose that CS-GAGs act as an endogenous, developmentally available ligand class that tunes microglial state, and that the balance between GAG deposition and lysosomal catabolism determines whether microglia return to homeostasis or persist in a phagocytic, DAM/PAM-like configuration.

Collectively, our findings identify lysosomal GAG catabolism as a fundamental regulator of microglial homeostasis and establish disrupted GAG turnover as a mechanistic link between extracellular matrix remodeling, lysosomal dysfunction, and disease-associated microglial activation. These results provide a framework for understanding how defects in ECM catabolism contribute not only to MPS VII, but also to broader neurodevelopmental and neurodegenerative disorders characterized by lysosomal stress and altered microglial function.

## ACKNOWLEDGEMENTS

We thank Dhruv Balaji, Crystal Tan, Marie-Anne Tomaj, Shaivam Kale, Neelanjana Komanduri, Shirya Parachuri, Jothsna Sabbasani, and Ronit Pullemla for assistance with data analysis. We thank Komlan Atitey for assistance with bioinformatics. We thank Paul Pavicic and Immunomonitoring Lab from Cleveland Clinic for processing PARSE single cell samples. We thank Brian Davis from PARSE Biosciences for assistance with protocols for cell fixation for PARSE scRNA-seq. We also thank Drs. Catherine Collins, Sam Pappas, Ranjan Dutta, Evan Deneris, and Peng Zhang for their critical review of the manuscript, and the staff of the Case Western Reserve University Animal Resource Center and Applied Functional Genomics Core for technical assistance. We thank Geert-Jan Boons (CCRC) for synthesizing the G0S0 NRE standard. Schematics were generated using Biorender (https://BioRender.com). This work was supported in part by the National Institute of Neurological Disorders and Stroke (R01NS122990 and R21NS144683 to D.Y.; R01NS122990-04S1 to KH) and the Mizutani Foundation for Glycoscience (grant 240071 to D.Y.). R.J.W. is supported by grants from NIH (R21NS135412) and the Cure Sanfilippo Foundation. The carbohydrate analyses performed at the CCRC were supported by NIH grant R24GM137782.

## AUTHOR CONTRIBUTIONS

K.H. and D.Y. designed research; K.H., B.L., Y.Z., A.B., K.C., and D.Y. conducted experiments; K.H., D.B., and C.T. performed image analyses; K.H. and D.Y. generated primary microglial cultures and performed phagocytosis assays; isolated and processed MACS microglia for scRNA-seq; D.Y and K.H performed scRNA-seq analyses with input from K.A; K.H. and Y.Z. performed β-glucuronidase activity assays; K.H. and Y.Z. performed mRNA isolation and qRT-PCR; R.W. and A.B. performed and analyzed HILIC-Q-TOF-MS of GAGs; Y.Z. performed RNAscope imaging and analysis; K.H., D.Y., R.W., A.B., and Y.Z. performed statistical analysis; and K.H. and D.Y. wrote the manuscript with input from all authors.

## DATA AVAILABILITY

The accession numbers for the datasets used for reporting; DAM scRNAseq expression (GSE98971); Proliferative region-associated microglia (GSE123030); Microglia Single Cell Atlas (GSE121654) (https://www.microgliasinglecell.com). Microglial scRNAseq dataset generated for this study will be deposited upon submission in Gene Expression Omnibus. All further study data is represented either within this manuscript or within supporting information. Further inquiries can be directed to the corresponding author.

## METHODS

### Generation and Maintenance of Mice

Procedures and mouse maintenance were performed in accordance with protocols approved by the Institutional Animal Care and Use Committee (IACUC) of Case Western Reserve University (CWRU). *Gusb^mps^* mice (B6.C-*H2-K^bm^*^1^/ByBir-*Gusb^mps^*/J JAX:000256) and E2a-Cre (B6.FVB-Tg(EIIa-cre)C5379Lmgd/J; JAX:003724) were purchased from Jackson Laboratory. *Gusb* floxed mice were generated in collaboration with Gempharmatech. To produce *Gusb^-/-^*mice, we crossed *Gusb^flx/flx^* mice with E2A-cre mice (germline Ella-Cre deletor; JAX:003724), generating *Gusb^+/-^* (GUSB-HET) mice, which we then interbred to create *Gusb^-/-^* (GUSB-KO) mice. Mice were genotyped using the primers GusKO_F (CTGTTGGCCTTCTTCCTTGTA), GusKO_R (AAGCTTCGGTTGCCTCTATG), and GUS-FLX-R (TGGCTGCATCCACAAGTGAAC), producing PCR band sizes of 192bp (WT allele), 297bp (FLX allele) and 372bp (KO or deletion of FLX allele).

### Histology and immunohistochemistry

Mice were deeply anesthetized with ketamine/xylazine (100/10 mg/kg, i.p.) or 20% urethane (i.p.) and transcardially perfused with 0.01 M PBS followed by 4% PFA in PBS. Brains were post-fixed overnight at 4°C, cryoprotected in 30% sucrose for 48 h, and cryosectioned coronally at 40 µm for free-floating or 16 µm for slide-mounted IHC. Free-floating sections were stored in PBS containing 0.1% sodium azide. Sections were washed in PBS, blocked for 1 h at room temperature in 5% normal donkey serum and 0.3% Triton X-100 in PBS, and incubated with primary antibodies for 48 h and secondary antibodies overnight, both at 4°C, with intervening PBS washes. Slide-mounted sections were processed similarly, with primary incubation overnight at 4°C and secondary incubation for 4 h at room temperature. Antibodies are listed in Table S5. For reconstruction of microglial morphology, rabbit anti-IBA1 and anti-TMEM119 were both used as the primary in the same channel to improve image quality.

#### Image Acquisition

Sections were imaged on a Zeiss Axio Imager Z1 equipped with an ApoTome.3 or on an Olympus VS200 slide scanner. High resolution images were acquired with 40x or 63x objectives using a Zeiss LSM 800 confocal microscope.

#### Image Analyses

Fiji was used to quantify the densities of WFA⁺ accumulations, enlarged CD68⁺ lysosomes, CLEC7A⁺ microglia and total microglia. These were measured in the corpus callosum (CC) or 300–500 µm-wide radial strips of primary somatosensory cortex (SSp) spanning layers 1–6. Lysosomes (LAMP1 or CD68) were reconstructed in three dimensions using Imaris (Oxford Instruments), and lysosome density and volume were quantified from the reconstructions. For microglia morphology, soma volume was measured in 40 µm sections. Branch number and Sholl analysis were measured in 300 µm sections, which preserve complete arbors. Individual microglia were reconstructed in Imaris to quantify soma volume and branch number, and Sholl analysis was performed with the Imaris Filament Sholl module.

#### Cortical dissociation

Mice were anesthetized with 20% urethane and transcardially perfused with 0.1 M PBS. Brains were removed, and meninges were cleared by rolling the tissue on sterile Whatman filter paper. Cortices were microdissected, gently triturated in 1 ml Accutase/Accumax (Sigma), and digested in 5 ml papain dissociation solution (5 mL EBSS supplemented with 0.5 mM EDTA, 3 mg/mL cysteine, and 1 mg/mL papain) pre-warmed to 37°C for 30 min. Digestion proceeded at 37°C for 25–30 min, with trituration once at 15–20 min. Cells were pelleted, resuspended in 5 ml 25% stock isotonic Percoll (SIP), and centrifuged at 600 × g for 20 min to remove myelin and debris. The upper layer and supernatant were discarded, and cells were washed in 10 ml wash buffer (1xHBSS/10%FBS). The resulting suspension was used for MACS isolation of microglia or for primary glial culture.

#### Magnetic-activated cell sorting (MACS) of microglia

Dissociated cells were resuspended in 1 ml MACS buffer (PBS with 0.5% BSA and 2 mM EDTA), pelleted, and resuspended in 50 µl buffer containing 5 µl CD11b MicroBeads (Miltenyi Biotec). After incubation for 15 min at 4°C, cells were washed in 1 ml buffer, resuspended in 500 µl, and applied to pre-equilibrated MS columns in a MACS separator. The unbound flow-through, the non-microglial (NMG) fraction, was collected. Columns were washed six times with buffer and removed from the magnet. Retained CD11b⁺ cells (microglia, MG) were eluted with 2 × 750 µl buffer using the column plunger. MG and NMG fractions were pelleted and stored at −80°C until analysis.

#### Primary microglial culture

Dissociated cortical cells from P5–P6 pups were washed in a 1:1 mixture of wash buffer and mixed glial culture medium (DMEM/F12/AA/10% FBS), resuspended in mixed glial culture medium, and plated in T75 flasks. Cultures were maintained at 37°C in 5% CO₂, with half of the medium replaced every 2 days for 10–14 days, until microglia proliferated on a confluent astrocyte monolayer. Microglia were then isolated by shaking for 1–2 hours on a rotary shaker inside the incubator, as previously described^75^.

#### Phagocytosis assay

Primary microglia, isolated as described above, were seeded onto glass coverslips in 24-well plates in microglial culture medium^75^. Cells were treated overnight with LPS (100 ng/mL), CSPG (100 µg/mL), LPS + CSPG, or vehicle control. All incubations were performed at 37 °C in 5% CO₂. Myelin conjugated to Alexa Fluor 647 NHS ester (Lumiprobe, #16820) was then added to each well at 200 µg/mL and incubated for 6 h. Acridine orange (1:500; Thermo Scientific, #6130) was added for the final 30 min of incubation to label acidic compartments. Coverslips were then washed with PBS, fixed in 4% PFA for 15 min, washed, and counterstained with DAPI (1:1000) in 5% normal donkey serum for 30 min at room temperature. After a final wash, coverslips were mounted onto slides with mounting medium for imaging.

#### β-Glucuronidase activity assay

The assay was conducted as previously described^76^. Briefly, tissue or cell pellets were lysed in 0.2 M sodium acetate buffer (pH 4.5) containing 10 mM EDTA and 0.1% Triton X-100. Serially diluted lysates (50 µl) were incubated with 50 µl of 10 mM 4-methylumbelliferyl β-D-glucuronide (MUG; Sigma) in 0.2 M sodium acetate (pH 4.5) for 1 h at 37°C. Reactions were stopped with 100 µl of 0.2 M Sodium Bicarbonate. Fluorescence (excitation 360 nm, emission 460 nm) was measured on a Spectramax microplate reader. Activity was calculated from a standard curve of purified β-glucuronidase (Sigma) and normalized to total protein using BCA.

#### RNA extraction and qRT-PCR

Total RNA extraction for qRT-PCR analysis was done using Trizol (Thermofisher) for mouse cerebral cortices per manufacturer’s instructions. cDNA synthesis from total RNA was done using MMLV Reverse Transcriptase (Takara) as per manufacturer’s instructions. Quantitative real time PCR (qRT–PCR) was performed with the StepOnePlus System (ABI) and 2x SYBR Power Mix (ABI) using QuantStudio 3 Real-time PCR system software. Primer sequences are listed in table S6.

### GAG isolation and disaccharide analysis

Glycosaminoglycans (GAGs) were isolated from MACS-enriched microglia (MG) and non-microglial (NMG) fractions from P14 cortex for quantitative disaccharide and non-reducing end (NRE) analysis, using previously reported methods^57^. Briefly, cell pellets were lysed using 0.5% CHAPS lysis buffer (50 mM HEPES, 120 mM NaCl, 2 mM EDTA, pH 7.4) containing a protease inhibitor cocktail (Roche). 50 µl of cell lysate was set aside for protein quantification via BCA assay. Next, cell lysates were diluted 1:10 in wash buffer (50 mM sodium acetate, 200 mM NaCl, pH 6.0) containing 0.1% Triton X-100 and Pronase (0.4 mg/mL; Sigma-Aldrich). Samples were digested overnight at 37°C with gentle agitation. Following digestion, samples were centrifuged (4,000 x*g*, 20 minutes), and the clarified supernatants were applied to DEAE-Sephacel columns (Cytiva) equilibrated with 50 mM sodium acetate buffer, pH 6.0, containing 200 mM NaCl. The resulting eluates were desalted using PD-10 desalting columns (Cytiva) and lyophilized prior to storage at -20°C.

For HS disaccharide analysis, purified GAGs were digested with heparin lyases I, II, and III (2 mU each; IBEX) for 16 h at 37°C in 40 mM ammonium acetate containing 3.3 mM calcium acetate, pH 7.0. For CS disaccharide analysis, purified GAGs were digested with chondroitinase ABC (20 mU, Sigma, # C3667) for 16 h at 37°C in a buffer containing 50 mM Tris and 50 mM NaCl, pH 7.9. The enzymatically depolymerized HS and CS disaccharides were labeled by reductive amination with (^12^C_6_) aniline (Sigma), as previously described^77^. Commercial HS and CS disaccharide standards (Iduron), labeled with (^13^C_6_) aniline (Sigma), were added to each sample as internal standards for absolute quantification of disaccharides.

HS and CS disaccharides were quantified by hydrophilic interaction liquid chromatography mass spectrometry (HILIC-LC/MS) using a Waters SYNAPT XS mass spectrometer coupled to an ACQUITY UHPLC H-Class system equipped with a BEH Glycan column (2.1 x 150 mm, 1.7 μm). Mobile phase A consisted of 50 mM ammonium formate buffer (pH 4.4, Waters), and mobile phase B consisted of 100% acetonitrile. Chromatographic separation was performed using 10% mobile phase A under isocratic conditions from 0 to 5.0 min, followed by a linear increase from 10% to 33% A from 5.0 to 49.0 min and from 33% to 45% A from 49.0 to 51.5 min. The column was maintained at 45% A from 52.0 to 60.0 min and subsequently washed with 90% B for 10 min. The column was maintained at room temperature, with a flow rate of 0.5 mL/min and an injection volume of 2 µL. Cumulative extracted ion chromatograms were generated, and data were processed using MassLynx software (Waters Corporation). HS and CS disaccharide abundance were quantified by comparison with the corresponding (^13^C_6_) aniline-labeled internal standards. Total HS and CS levels were normalized to protein amount measured via BCA assay (Thermo).

### Non-reducing end (NRE) analysis

GAGs isolated from MACS-enriched microglia (MG) and non-microglial (NMG) fractions from P14 cortex were used for NRE analyses. GAGs were isolated, as described above, and were subsequently enzymatically depolymerized with heparin lyases or chondroitinase ABC and were differentially mass labeled by reductive amination with (^12^C_6_) aniline, as described above. HS NRE species (G0S0) were analyzed by mixing digested HS with (^13^C_6_) aniline-tagged β-D-glucopyranosyluronate-(1-4)-*O*-2-deoxy-2-*N*-sulfamino-α/β-D-glucopyranose (G0S0, 10 pmol). The G0S0 disaccharide standard was prepared as previously described^57^. Samples were analyzed by HILIC-LC/MS, as described above. G0S0 NRE species were quantified using the isotopically labeled internal standard and normalizing to total protein, as measured by BCA.

Loss of β-glucuronidase is predicted to leave a terminal β-GlcA at the non-reducing end of CS chains, which chondroitinase ABC releases as a saturated disaccharide, GlcA-GalNAc(4S or 6S) (G0a4/G0a6). Because authentic standards for these species are not available, G0a4 and G0a6 were not resolved and are reported together as G0a4/G0a6. This species was identified by accurate mass of the (^12^C_6_) aniline-tagged (M-H⁻) ion (theoretical m/z 553.135; obs m/z 553.133), consistent with previous reports^57^. G0a4/G0a6 abundance was determined from extracted ion chromatogram peak area and normalized to total protein.

#### Single Cell RNAseq

Single-cell suspensions of MACS purified microglia were fixed and permeabilized using the Evercode Cell Fixation kit (Parse Biosciences) according to the manufacturer’s instructions. Fixed cells were counted and processed using the Evercode Whole Transcriptome Mini v3 kit at CCF Immunomonitoring lab. RNA was reverse-transcribed, barcoded and sequencing libraries generated at Immunomonitoring lab (CCF) for sequencing. After initial sequencing, quality-control filtering removed low-quality cells (from high doublet score (based on Trailmaker Doublet scores)) and high-mitochondrial cells (>15% mitochondrial transcripts) yielding high-quality transcriptomes for downstream analysis. Highly variable genes were identified using the variance stabilizing transformation. PCA was performed using the top 2,000 HVG (Highly Variable Genes). Following normalization and PCA, datasets were integrated using the Harmony algorithm to correct for batch effects while preserving biological variation. The first 30 principal components were used for graph construction and UMAP visualization. Clustering was performed using the Leiden algorithm (resolution = 0.1 and Minimum Distance = 0.1).

#### RNAscope

Single-molecule fluorescent in situ hybridization was performed using the RNAscope Multiplex Fluorescent Reagent Kit v2 (Advanced Cell Diagnostics (ACD), Newark, CA) according to the manufacturer’s instructions. Mice were transcardially perfused with PBS followed by 4% paraformaldehyde; brains were post-fixed overnight in 4% PFA at 4 °C, cryoprotected in 30% sucrose, embedded in OCT, and cryosectioned at 16 µm. Frozen sections were mounted on SuperFrost Plus slides and stored at −80 °C until use. Sections were pretreated with hydrogen peroxide, target retrieval, and protease per the ACD protocol. Sections were then hybridized with target probes against Apoe (Cat. No. 313271-C2, channel C2) and *Tmem119* (Cat. No. 472901-C3, C3), for 2 h at 40 °C in a HybEZ oven, followed by sequential amplification (AMP1–3) and channel development with Opal dyes. Nuclei were counterstained with DAPI, and slides were mounted for imaging and analyses.

### Statistics

All graphed data is reported as mean ± SEM. All statistical tests reported (Student’s t-tests, <u>Chi-square tests</u>, One-way and two-way ANOVAs) were performed using GraphPad Prism software (version 11).

