## Supplemental Figures for "Lysosomal Glycosaminoglycan Catabolism Regulates Microglial Cell State in the Developing CNS"

### Supplemental Fig. 1

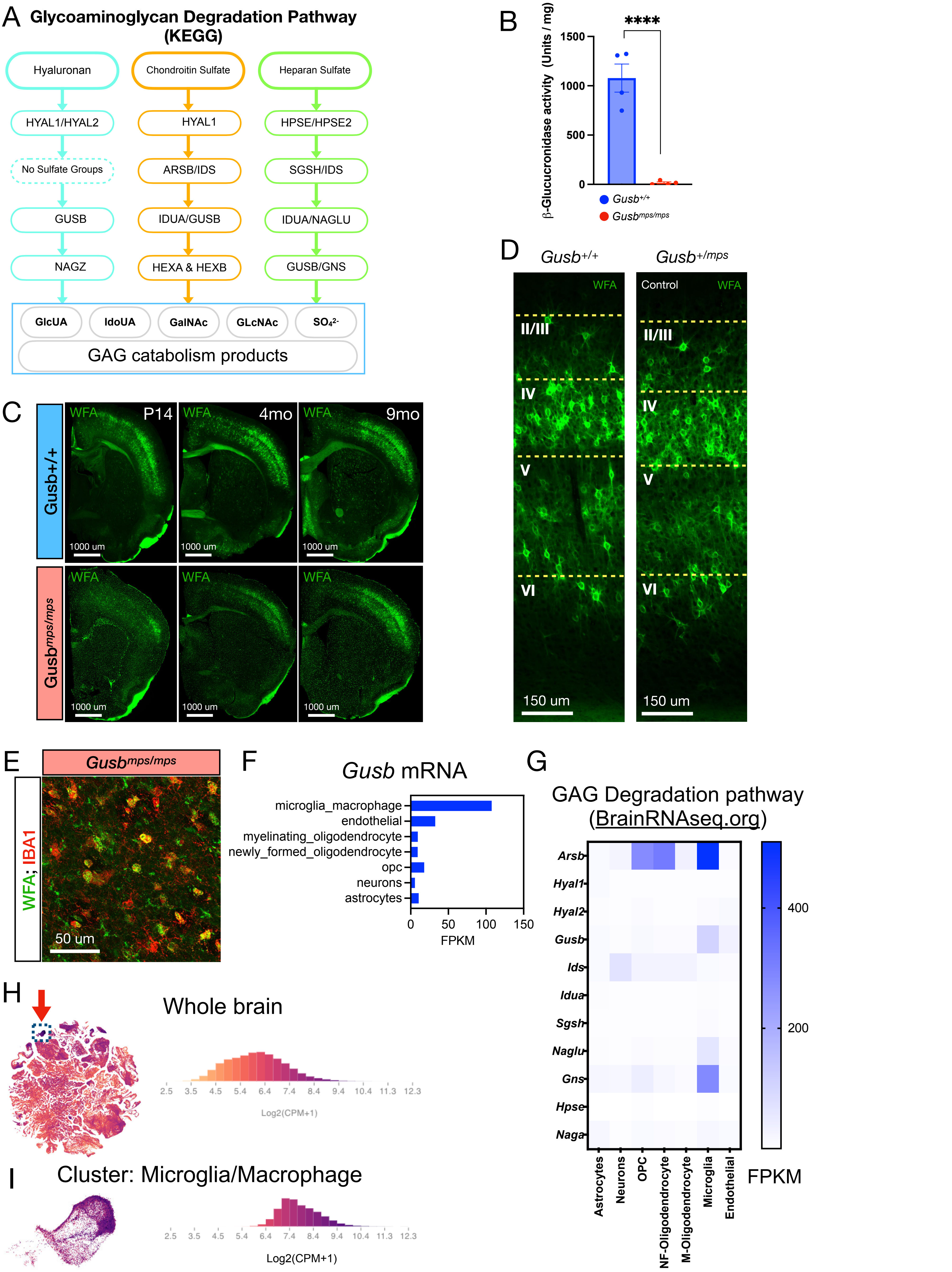

**Figure S1. Ablation of β-glucuronidase results in intracellular GAG aggregates in microglia.**
(A) Schematic (based on the KEGG pathway) depicting the enzymes involved in the degradation of chondroitin sulfate, heparan sulfate, and hyaluronic acid GAGs, with some of the resulting byproducts that result from the different processes. The role of β-glucuronidase in the pathway is highlighted. (B) Graph depicting the complete loss of β-glucuronidase activity (y-axis; β-glucuronidase activity/mg) in cortical tissue from *Gusb*$^{mps/mps}$ mice relative to *Gusb*$^{+/+}$ controls (x-axis; mouse genotype) as measured with enzyme assay (Welch's *t*-test, $t(3.075) = 18.52$, $P = 0.0003$, $N = 4$). (C) Representative images of hemisections from P14 (left), 4 mo (middle), and 9 mo (right) *Gusb*$^{+/+}$ and *Gusb*$^{mps/mps}$ mice stained for WFA (green) using IHC, demonstrating persistent WFA-positive GAG aggregates throughout the CNS of *Gusb*$^{mps/mps}$ mice that persist with aging. (D) Representative images of the SSp (primary somatosensory cortex) from P14 control (*Gusb*$^{+/+}$) and MPS-Het (*Gusb*$^{+/mps}$) mice spanning all cortical layers and stained for WFA (green). (E) Representative subcortical image from a Gusb$^{mps/mps}$ mouse stained for IBA1 and WFA demonstrating how the wfa accumulations colocalize with microglia throughout the CNS. (F) Graph of *Gusb* FPKM values (x-axis; FPKM of *Gusb*) for different cell types (y-axis; cell types) derived from the Brain RNA-seq database. Heatmap of the FPKM values of 16 GAG degradation pathway genes (y-axis) for different cell types (x-axis; cell types) derived from the Brain RNA-seq database. (H–I) UMAP plots and corresponding graphs showing *Gusb* expression abundance (x-axis, $log_2$CPM; y-axis, number of cells ×10,000) from 10x scRNA-seq data of (F) whole brain and (G) the microglial cluster derived from the ABC Whole Mouse Brain Atlas (Allen Brain Atlas).

### Supplemental Figure 2

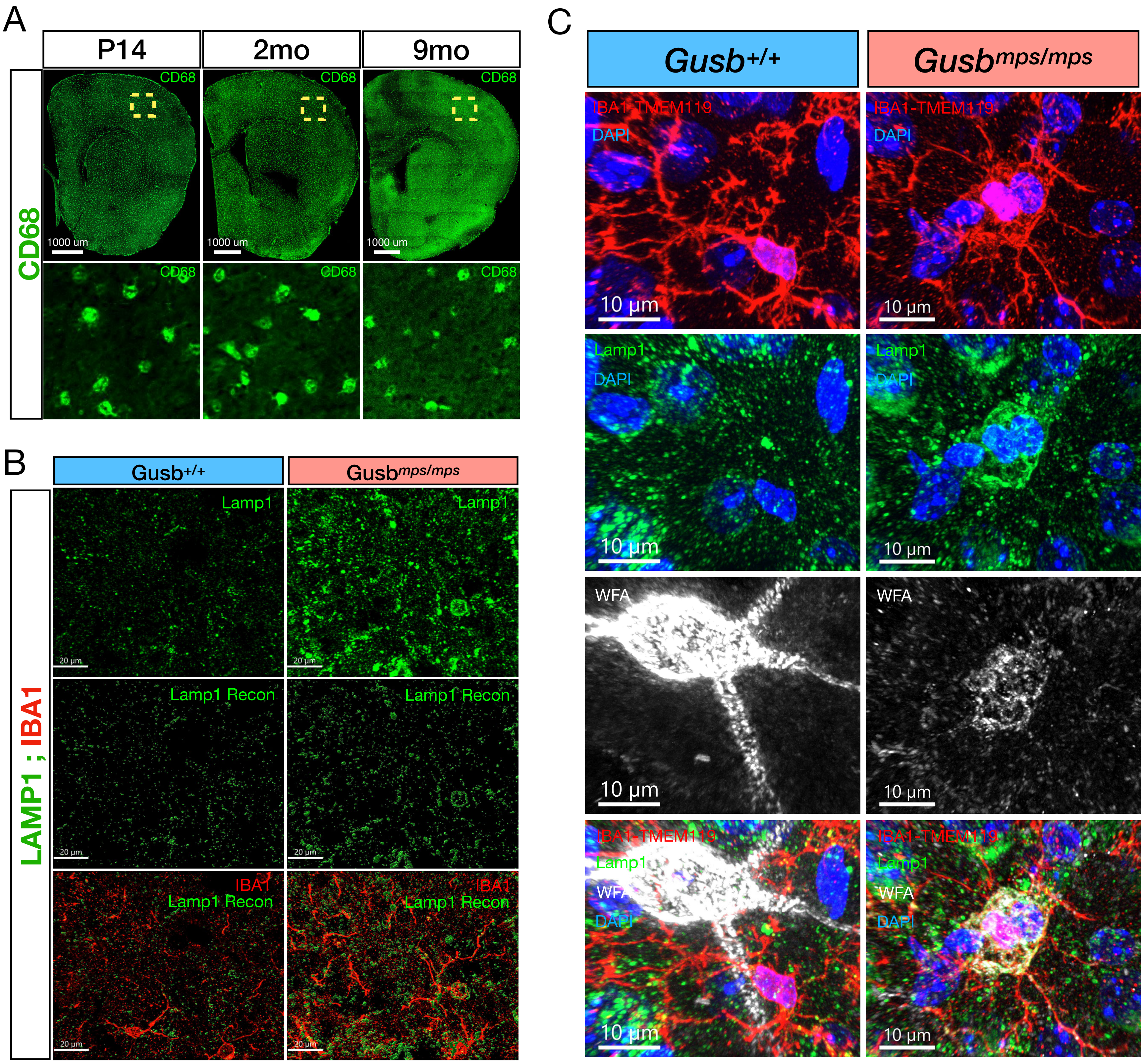


**Figure S2. Abnormal lysosomes containing WFA+ aggregates are observed across the CNS.**
(A) Representative images of hemisections from P14 (left), 4 mo (middle), and 9 mo (right) *Gusb*$^{+/+}$ and *Gusb*$^{mps/mps}$ mice stained for CD68 (green), demonstrating abnormal CD68-positive GAG-containing lysosomes throughout the entire hemisphere of *Gusb*$^{mps/mps}$ mice that persist with aging. (B) Representative images of SSp layer 4 from from *Gusb*$^{+/+}$ and *Gusb*$^{mps/mps}$ mice stained for DAPI and IBA1 (top row), LAMP1 (second row), WFA (third row), and the merged image (fourth row), demonstrating that enlarged LAMP1-positive lysosomes contain WFA-positive aggregates. (C) Representative images of LAMP1+ lysosomes in layer IV of the SSp colocalizing with WFA aggregates in *Gusb*$^{mps/mps}$ microglia. The tissue is strained for LAMP1, WFA, IBA1-TMEM119 and DAPI.

### Supplemental Figure 3

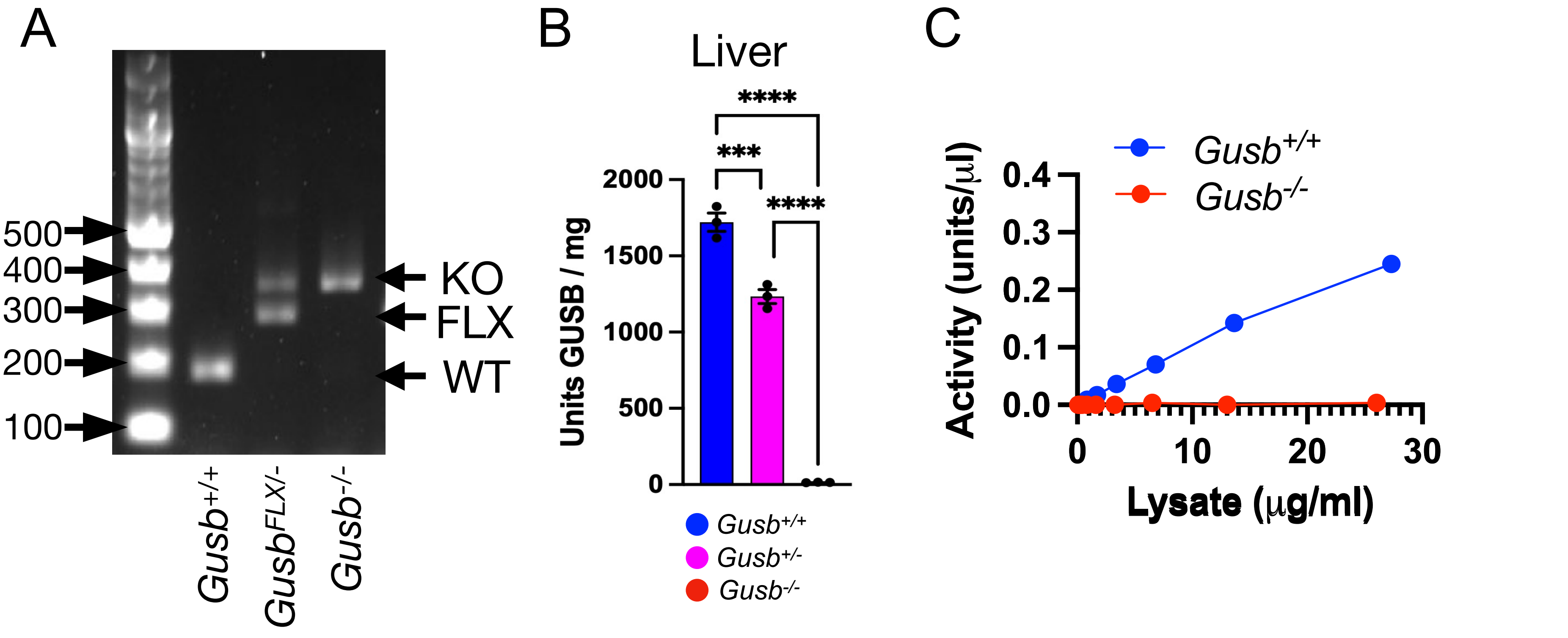


D

| Genotype frequencies of progeny from $Gusb^{+/-}$ X $Gusb^{+/-}$ cross | | | |
|---|---|---|---|
| Genotype | Observe | Ratio | Ratio (Expected) |
| $Gusb^{+/+}$ | 28 | 29.17 | 25 |
| $Gusb^{+/-}$ | 51 | 53.13 | 50 |
| $Gusb^{-/-}$ | 17 | 17.71 | 25 |

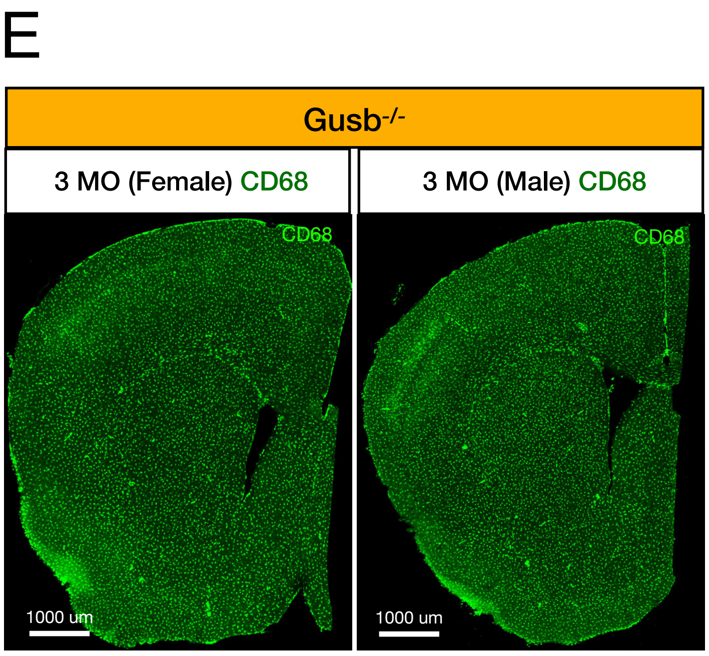


**Figure S3. Total loss of β-glucuronidase activity in *Gusb* null mice.**
(A) Image depicting genotyping results for the ***Gusb*** allele, demonstrating bands corresponding to WT ($Gusb^{+/+}$), Heterozygous($Gusb^{+/-}$) and null ($Gusb^{-/-}$) alleles. (B) Graph of β-glucuronidase activity in liver tissue from $Gusb^{+/+}$, $Gusb^{-/+}$, and $Gusb^{-/-}$ mice (one-way ANOVA, $F(2,6) = 414.2$, $P < 0.0001$, Dunnett's multiple comparisons test with Tukey correction, $N = 3$). (C) Graph of β-glucuronidase activity (y-axis; β-glucuronidase activity/mg) in cortical tissue from $Gusb^{+/+}$ and $Gusb^{-/-}$ mice (x-axis; mouse genotype) measured across increasing concentrations of tissue lysate (x-axis; lysate concentration ug/ml). (D) Table showing expected and observed genotype frequencies of progeny from $Gusb^{+/+}$ and $Gusb^{-/-}$ crosses. (E) CD68 stained hemisections from 3 month old male and female $Gusb^{-/-}$ mice depicting no changes in the

### Supplementary Figure 4

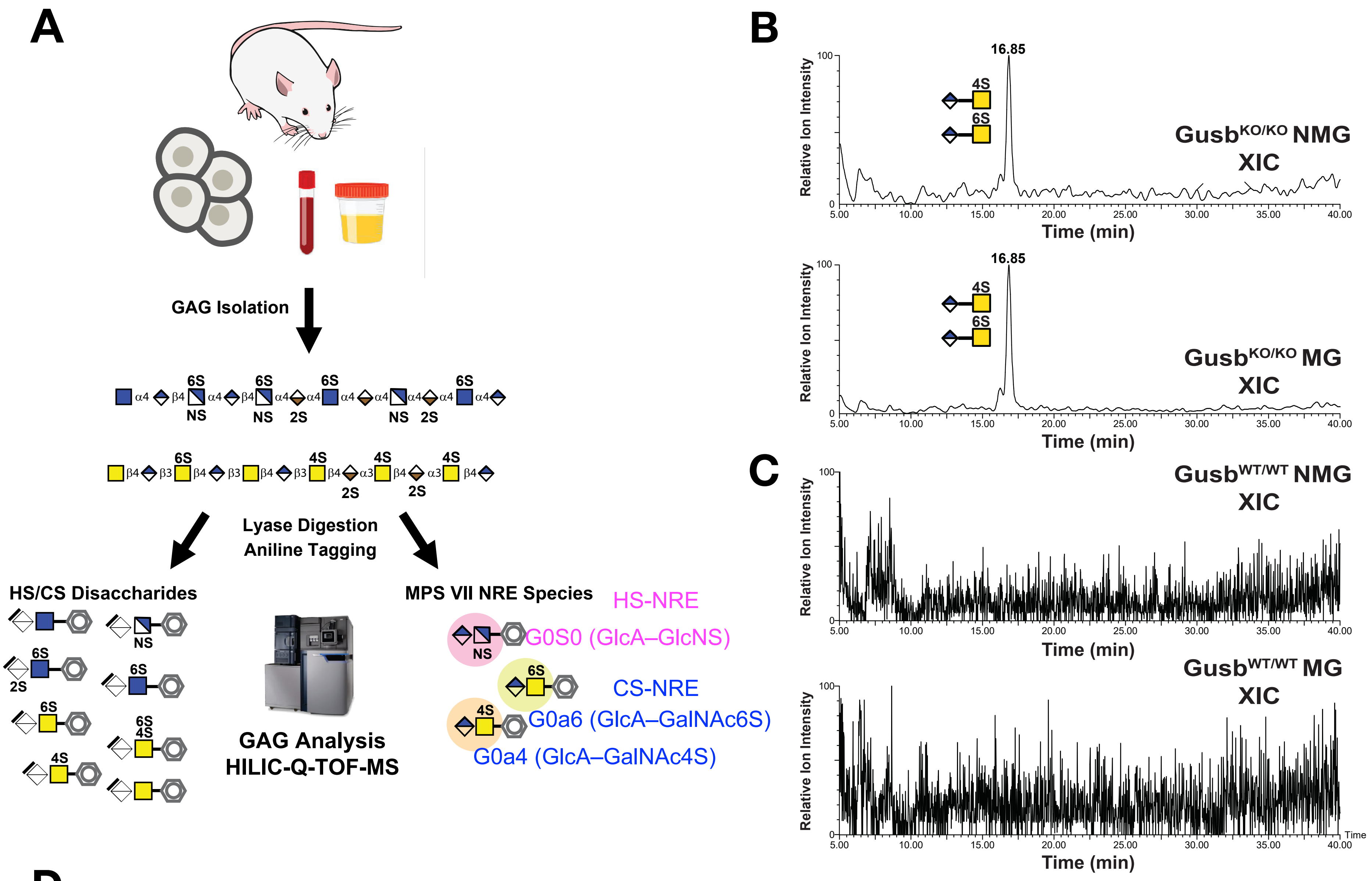


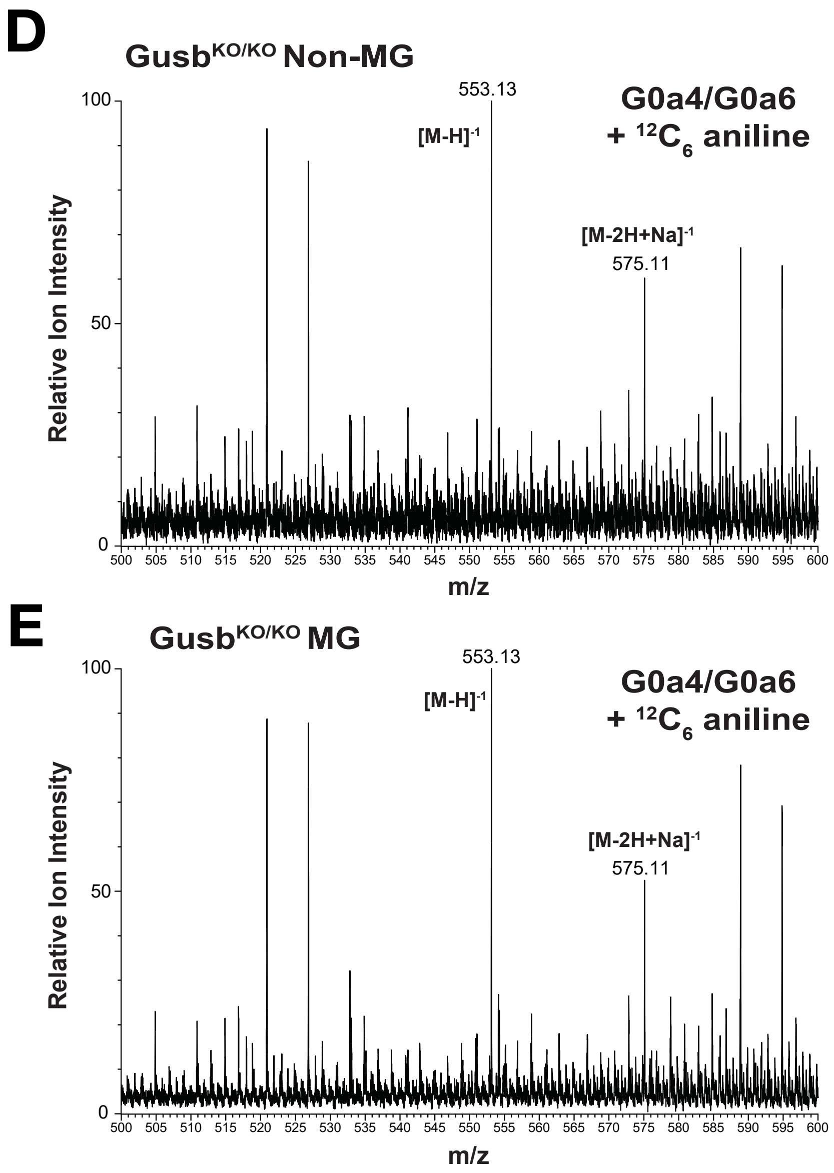


**Figure S4. Detection of chondroitin sulfate non-reducing end species in *Gusb*-deficient cells.** (A) Schematic depicting the analyses of total and NRE GAGs by disaccharide analyses. (B, C) Extracted ion chromatograms (m/z 553.13) of [$^{12}C_6$]aniline-tagged chondroitinase ABC digests from MACS-enriched microglia (MG) and non-microglial cells (NMG) of P14 (B) *Gusb*$^{-/-}$ and (C) *Gusb*$^{+/+}$ cortex. The minor earlier-eluting component (~16.3 min) may represent the alternate sulfate positional isomer. (D, E) Mass spectra across the 16.85 min peak in P14 (D) *Gusb*$^{-/-}$ and (E) *Gusb*$^{+/+}$ cortex. showing the $[M{-}H]^-$ (m/z 553.13) and $[M{-}2H{+}Na]^-$ (m/z 575.11) ions of the tagged NRE species (G0a4/G0a6). Because authentic standards are unavailable, 4-*O*- and 6-*O*-sulfated isomers (G0a4/G0a6) were not distinguished. Traces are representative of n =3 per group.

### Supplemental Figure 5

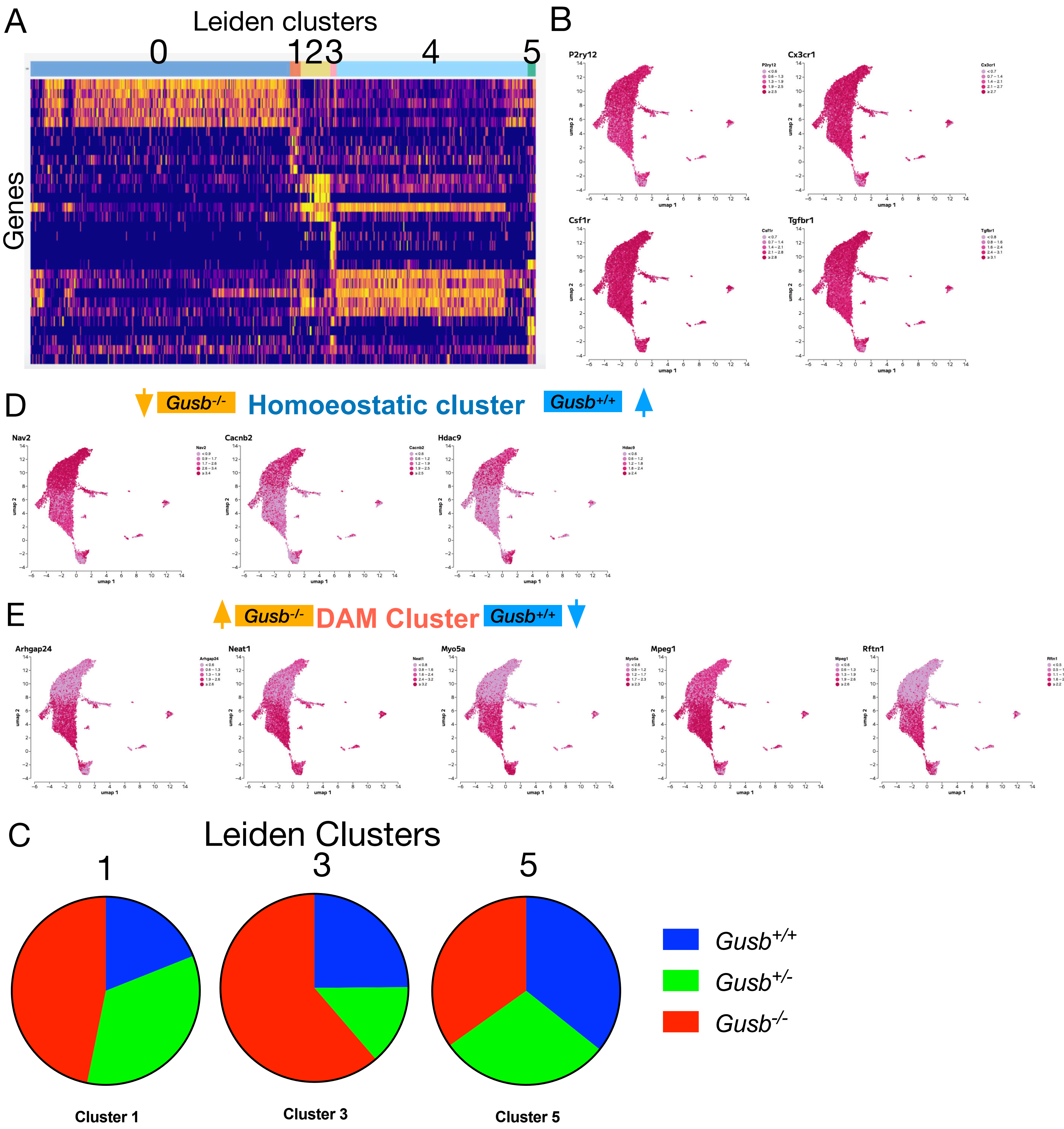


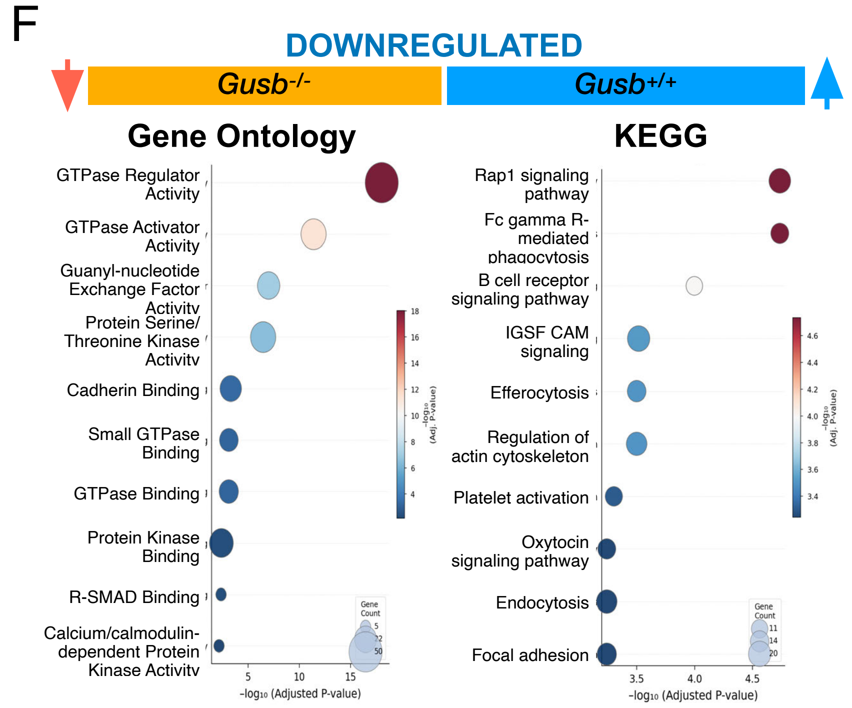


**Figure S5. *Gusb* deficient microglia are transformed into a phagocytic DAM like cell state.** (A) Heatmap depicting relative expression of DEGs across the leiden clusters. (B) UMAP plot depicting the expression of canonical microglial lineage genes in across the leiden clusters. (C-D) UMAP plot depicting the expression of candidate (C) downregulated and (D) upregulated DEGs in *Gusb*$^{-/-}$ relative to *Gusb*$^{+/+}$ across the leiden clusters. (E) Pie chart depicting the relative proportion of (blue) *Gusb*$^{+/+}$, (green) *Gusb*$^{-/+}$, and (red) *Gusb*$^{-/-}$ in leiden clusters 1, 3 and 5. (F) Bubble plot showing KEGG and Gene Ontology pathway enrichment among genes upregulated in *Gusb*$^{+/+}$ microglia relative to *Gusb*$^{+/+}$ controls. Bubble size indicates the number of differentially expressed genes associated with each pathway, and color intensity represents enrichment significance (false discovery rate-adjusted *P* value).

### Supplemental Figure 6

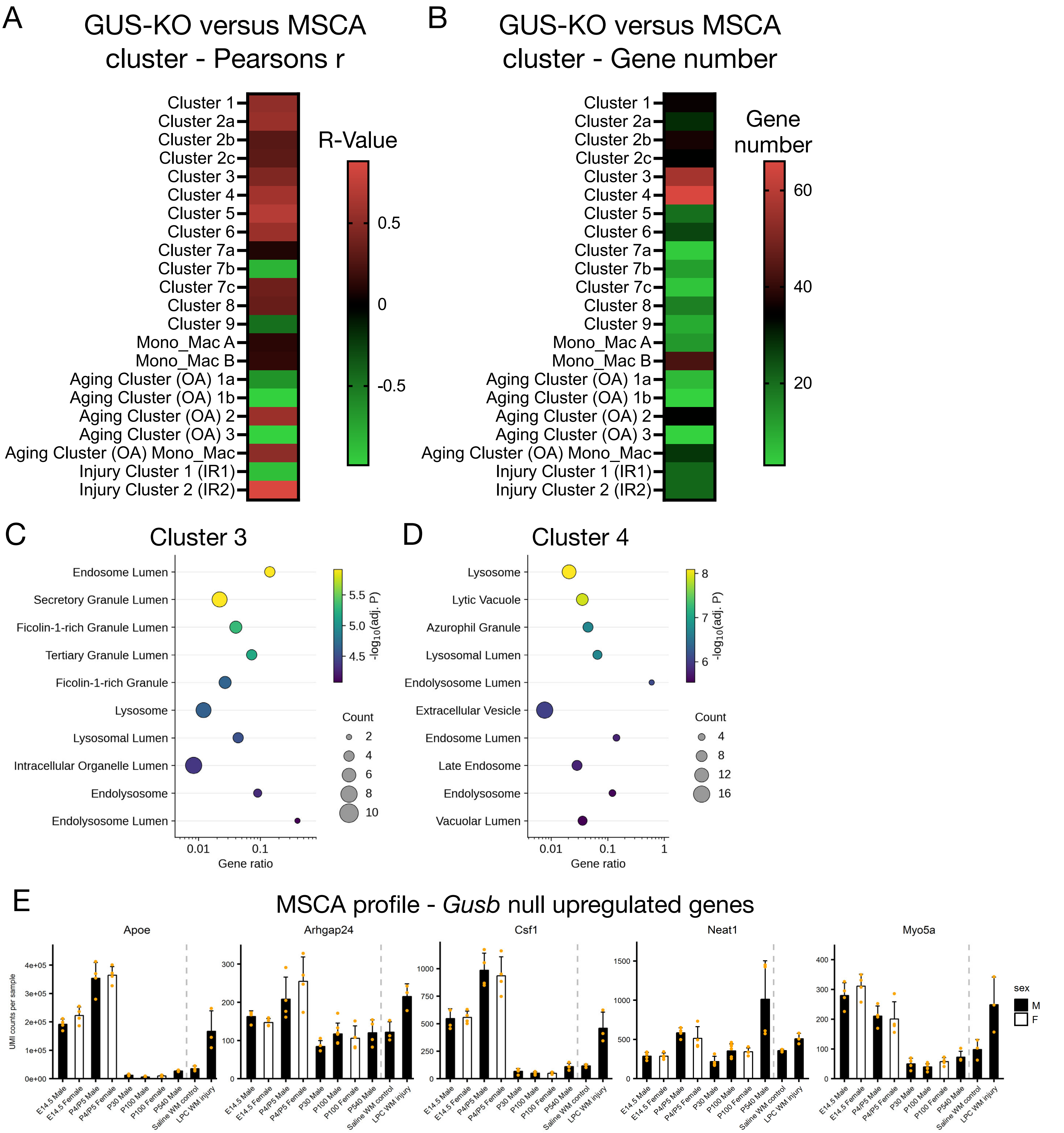


**Figure S6. *Gusb* deficient microglia are transcriptionally resemble naive microglia.**
(A-B) Heatmaps depicting the similarities of GUS-KO cluster microglia with the different Microglia Single Cell Atlas clusters, via (A) measurement of their coorelation strength via Pearson's r-value, and (B) the number the DEGs that they have in common, with developmental clusters 3 and 4 have both a notable high r-value and number of genes in common. (C-D) Bubble plots showing Cellular Ontology pathway enrichment among genes upregulated in Microglia Single Cell Atlas clusters 3 (C) and 4 (D). Bubble size indicates the number of differentially expressed genes associated with each pathway, and color intensity represents enrichment significance (false discovery rate-adjusted *P* value). (E) Bar graphs of the normal expression patterns in microglia from male and female mice across various time points for some of the top upregulated genes in *Gusb*$^{-/-}$ microglia relative to *Gusb*$^{+/+}$ controls (in order from left to right, top to bottom), genes downregulated in *Gusb*$^{-/-}$ microglia are highly expressed in early development. Graphs were generated using the Microglial Single Cell Database.

### Supplemental Figure. 7

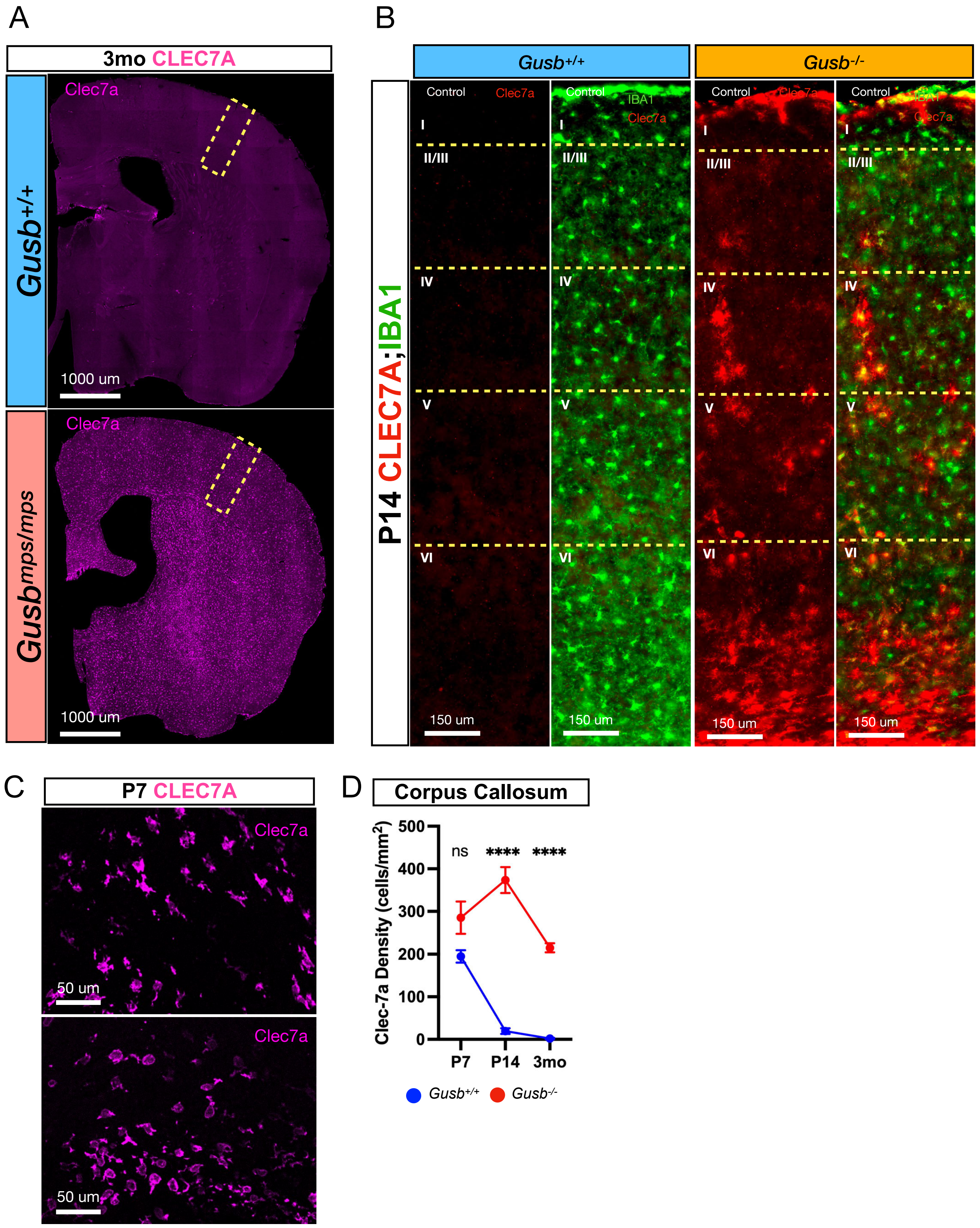


**Figure S7. Microglial expression of CLEC7A in GUSB-deficient mice.**
(A) Representative images of hemisections from 3mo Gusb$^{+/+}$ and *Gusb*$^{mps/mps}$ mice stained for CLEC7A (magenta), demonstrating robust expression of CLEC7A in *Gusb*$^{mps/mps}$ microglia. (B) Representative images of the P14 SSp from *Gusb*$^{+/+}$and *Gusb*$^{-/-}$ mice spanning all cortical layers and stained for CLEC7A (red) and IBA1-TMEM119 (green). (C-D) Representative images (C) and corresponded quantification (D) of the CC from P7 *Gusb*$^{-/-}$ and *Gusb*$^{+/+}$ mice immunostained for CLEC7a (magenta) demonstrating similar expression of CLEC7a in microglia from both *Gusb*$^{+/+}$ and *Gusb*$^{-/-}$ mice (Student's *t*-test: $t(6) = 2.216$, ns, N=2).

### Supplemental Figure 8

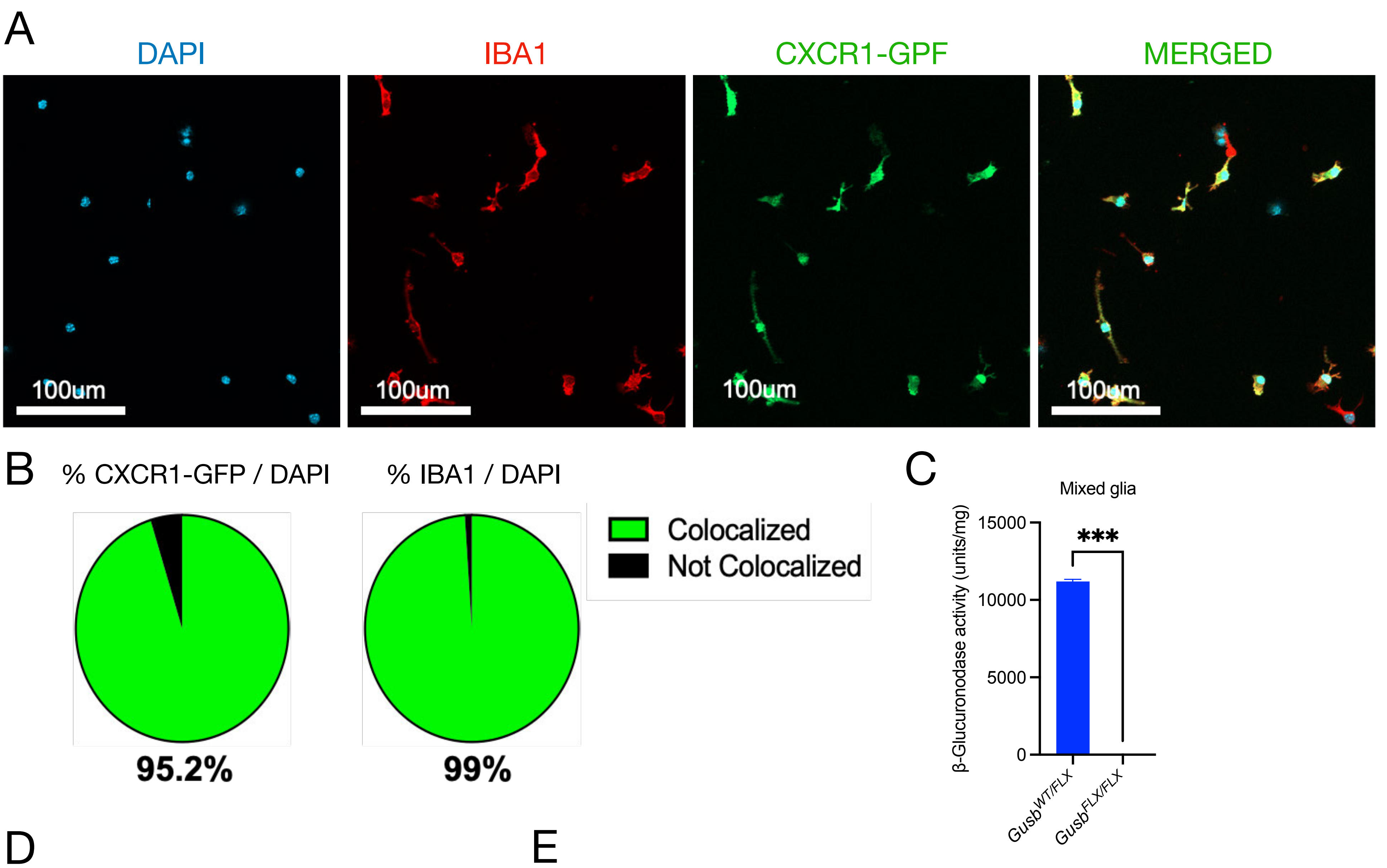


D

| | | Gusb+/+ Blank | Gusb+/+ CSPG | Gusb+/+ LPS | Gusb+/+ CSPG +LPS | Gusb-/- Blank | Gusb-/- CSPG | Gusb-/- LPS | Gusb-/- CSPG +LPS |
|---|---|---|---|---|---|---|---|---|---|
| *Gusb+/+* | Blank | ns | **** | **** | **** | **** | **** | **** | **** |
| | CSPG | **** | ns | ns | * | ns | **** | ns | **** |
| | LPS | **** | ns | ns | ns | ns | **** | ns | **** |
| | CSPG+ LPS | **** | * | ns | ns | ns | **** | ns | **** |
| *Gusb-/-* | Blank | **** | ns | ns | ns | ns | **** | ns | **** |
| | CSPG | **** | **** | **** | **** | **** | ns | **** | ns |
| | LPS | **** | ns | ns | ns | ns | **** | ns | **** |
| | CSPG+ LPS | **** | **** | **** | **** | **** | ns | **** | ns |

E

| | | Gusb+/+ Blank | Gusb+/+ CSPG | Gusb+/+ LPS | Gusb+/+ CSPG +LPS | Gusb-/- Blank | Gusb-/- CSPG | Gusb-/- LPS | Gusb-/- CSPG +LPS |
|---|---|---|---|---|---|---|---|---|---|
| *Gusb+/+* | Blank | ns | **** | ns | ns | *** | **** | **** | **** |
| | CSPG | **** | ns | ns | **** | ns | **** | ns | ns |
| | LPS | ns | ns | ns | ns | ns | **** | ns | **** |
| | CSPG+ LPS | ns | **** | ns | ns | ** | **** | **** | **** |
| *Gusb-/-* | Blank | *** | ns | ns | ** | ns | **** | ns | * |
| | CSPG | **** | **** | **** | **** | **** | ns | **** | *** |
| | LPS | **** | ns | ns | **** | ns | **** | ns | * |
| | CSPG+ LPS | **** | **** | **** | **** | * | *** | * | ns |

**Figure S8. Isolation of primary microglial cultures**
Representative images (A) and corresponding pie charts (B) of microglia isolated from $CX_3CR\text{-}1_{GFP}$ knock-in cortex and stained for DAPI (blue), IBA1 (red), and GFP (green), demonstrating the high purity of the isolated microglial cultures. C) Graph depicting fold change (y-axis) of *Gusb* mRNA expression in RNA isolated from $Gusb_{Flx/Flx}$ and $Gusb_{-/-}$ (x-axis) mixed glial cultures. (D-E) Tables depicting the statistical comparisons using Two-way ANOVA for (D) myelin phagocytosis and (E) AO intensity from all experimental conditions (with and without CSPG/LPS) for control and $Gusb_{-/-}$ microglia.
